# Testing the efficacy of artificial flowers as a novel attractant for automated pollinator monitoring

**DOI:** 10.64898/2026.01.23.698111

**Authors:** Abra Ash, Stephen Hallett, Claire Carvell, Leon Williams, Tom August

**Affiliations:** UK Centre for Ecology & Hydrology, MacLean Bldg, Benson Ln, Crowmarsh Gifford, Wallingford OX10 8BB; Cranfield University, College Rd, Wharley End, Bedford MK43 0AL

**Keywords:** Monitoring, insect pollinators, camera trap, attractant preference, artificial flowers

## Abstract

Insect camera traps are a rapidly developing technology, enabling automated monitoring of insects. However, little has been reported on improving the attractants used for daytime flying insects on such cameras. This study compares the attractiveness of novel, 3D printed, artificial flowers with traditional methods of attracting insects (e.g. pan traps and solid coloured paper squares). We hypothesised that artificial flowers would attract a higher abundance and diversity of insects compared to traditional attractants by more accurately mimicking flowers.

Additionally, we examined colour preference and average landing duration on the attractants. Artificial flowers, dry pan traps and paper squares, painted in yellow, white, or blue ultraviolet fluorescent paint, were filmed simultaneously to observe wild insect behavioural responses (landings and approaches). The results indicate overall preference for artificial flowers over the two traditional attractants when considering all insect groups together, and overall colour preferences for blue and yellow. When analysing insect groups separately, hoverflies preferred landing on artificial flowers over the other attractants. Bumblebees preferred approaching artificial flowers, and small insects preferred landing and approaching artificial flowers over the other attractants. ‘Other flies’ preferred landing on pan traps and paper over artificial flowers. Hoverflies, small insects, wasps, and solitary bees responded more to yellow than the other colours, while bumblebees responded more to blue.

Comparisons of landing durations revealed that hoverflies spent longer on the artificial flowers than paper. ‘Other flies’ spent longer on the pan traps and paper. These results show that artificial flowers could offer an efficient attractant for insect camera traps as they attracted a higher abundance of key pollinating insects (hoverflies and bumblebees), and do not have worse attraction rates for the other insect groups (excluding ‘other flies’).

## Introduction

There is increasing evidence worldwide for changes in insect biodiversity and abundance having occurred over recent decades (Outhwaite et al. 2020; Klink et al. 2020). Terrestrial insects have been decreasing in abundance over the last century with these changes largely associated with land-use and land-management drivers and climate change (Klink et al. 2020; Hailay Gebremariam 2024). Areas with historic climate warming and intensive agricultural land use are associated with an almost 50% reduction in insect species abundance and this loss is alarming for its direct impact on biodiversity and ecosystem stability (Outhwaite et al. 2022). Flower-visiting insects, specifically butterflies, wild bees, hoverflies, and moths, are included in this decline and play a crucial role in providing ecosystem services such as pollination and food security (Klein et al. 2007). Evidence for their decline has led to a need for long-term population monitoring to better understand the current state of insect pollinator biodiversity, the impact of declines in pollinators, and to develop interventions that could help to reverse this decline (Stout & Dicks 2022; European Parliament; the Council of the European Union 2024).

Traditionally, a significant amount of pollinator monitoring has been undertaken by citizen scientists through volunteer-led recording schemes collating species records, such as the Hoverfly Recording Scheme (HRS) and the United States-based Bumble Bee Atlas, or by structured monitoring surveys such as the UK Pollinator Monitoring Scheme (PoMS) (UK Pollinator Monitoring Scheme 2025) and European Butterfly Monitoring Scheme (eBMS) (Powney et al. 2019; Roy et al. 2020). In addition, volunteers, in combination with expert taxonomists, collect insects from primarily lethal attractants, such as pan traps with soapy water (O’Connor et al. 2019; Rondeau et al. 2023), or sticky traps (Geissmann et al. 2022). In the context of this paper, we define an ‘attractant’ as an item designed to attract pollinating insects with different features such as shape and colour.

Insect camera trap designs were first introduced in 2011 and have become more popular since the early 2020’s for remote monitoring of insects (Lortie et al. 2011; Droissart et al. 2021; Bjerge et al. 2021; Geissmann et al. 2022; Sittinger et al. 2024; Roy et al. 2024). Using these hardware systems in conjunction with new analytical tools (e.g. computer vision) has the potential to identify insects, and allow long term, consistent monitoring (Naqvi et al. 2022; van Klink et al. 2022). Advances have been made in the development of insect camera traps that either use artificial intelligence (AI) to identify differences on the background or take timelapse photos.

Furthermore, insect identification software has improved, using machine learning to identify insects. However, little has been reported on improving and standardising the attractants used for daytime flying insects (Droissart et al. 2021; Besson et al. 2022; Bjerge et al. 2022, 2023; van Klink et al. 2022; Sittinger et al. 2024). Current daytime automated insect camera traps use a mix of lethal (e.g. sticky paper) (Geissmann et al. 2022) and non-lethal (e.g. 2D flower shapes printed on paper or solid-coloured screens) attractants (DIOPSIS 2020; Sittinger et al. 2024).

To further develop the attractants used in automated insect monitoring systems, we propose using realistic-looking artificial flowers that have the possibility of attracting a wider diversity and abundance of pollinating insects than existing methods. Artificial flowers, first introduced as tools to aid behavioural insect research in 1876 by Felix Plateau, allow researchers to control every element of the flower, leading to discoveries that would be impossible without them (Burton 1877; Clements & Long 1923; Campos et al. 2015). These behavioural studies cover the broad range of insect floral trait preferences such as colour (Lunau et al. 2006), scent (Witjes & Eltz 2007), shape (Campos et al. 2015), as well as many others (Clarke et al. 2013; Harrap et al. 2020). Furthermore, the advent of modern 3D printing technology allows construction of long-lasting artificial flowers, making them an ideal attractant in long-term automated monitoring. While artificial flowers have been widely used to study behaviour and toxicology in pollinators, they have not yet been applied to support automated insect monitoring or compared to traditional attractants.

This paper reports on an experiment comparing the effectiveness of traditional methods (pan traps and sticky paper) in attracting insects in the wild with the novel method of using artificial flowers. Sticky traps, with references dating back to the 14^th^ century, are still a commonly used method to trap flying insects (Fay 2022). Purchased for domestic and industrial use to trap pest insects, sticky traps often have solid colours (i.e. yellow) that attract insects from a distance (Sampson et al. 2021). There is already an overlap in artificial flowers using sticky traps and sticky traps being used in automated monitoring (Jürgens et al. 2015; Geissmann et al. 2022). Pan traps, first introduced in 1951 by V. Moericke, are a widespread, common method for monitoring insect populations (Heathcote 1957; Vrdoljak & Samways 2012; St. Clair et al. 2020). They comprise of coloured bowls filled with soapy water that keep insects trapped when they land and/or fall into the bowl (Baum & Wallen 2011).

We hypothesise that artificial flowers will attract a higher abundance and diversity of insects compared to the traditional monitoring methods as they more closely resemble real flowers. In determining this there are three objectives. First, we directly compare insect behavioural responses (landing and approaching) with the artificial flowers to insect behavioural responses with dry pan traps and coloured paper (mimicking the appearance of sticky traps). Second, we determine whether colour has an impact on the insect behavioural responses with the attractants, knowing that some insect groups have innate colour preferences and to show that using previously researched colours will aid in the attraction of wild insects. Third, we compare the average time insects spent on each attractant to determine whether the artificial flowers could retain insects under a camera for a longer time-period than traditional attractants, giving greater opportunity for high quality image collection. Lastly, we suggest further steps for consideration to create more effective and standardised attractants for future automated pollinator monitoring.

## Methods

Three different attractants were designed and tested in our experiment: artificial flowers, dry pan traps (coloured bowls) and coloured paper (to mimic sticky traps). Each attractant was presented in three ultraviolet (UV) fluorescent colours (yellow, white, and blue) using similar painting and colour choice methodology to the UK PoMS (O’ Connor et al. 2019). These three colours were chosen as different insect groups are attracted to different colours (e.g. yellow for wasps and hoverflies, and blue for bees) (Acharya et al. 2021).

### Artificial Flowers

The artificial flowers were designed using the 3D modelling software, ZBrush (Maxon 2025). The flowers were designed to be radially symmetrical with five petals and stamen in the centre to resemble the generalist flower shape of a buttercup (*Ranunculus* species) as it is radially symmetrical and bowl-shaped. It has features that have been found to attract of a wide variety of insects, and is one of the most generalist flower shapes (Figure 1) (Kevan & Baker 1983; Olesen et al. 2007; Woodcock et al. 2014; Yoder et al. 2020; Roquer-Beni et al. 2022).

**Figure 1.**
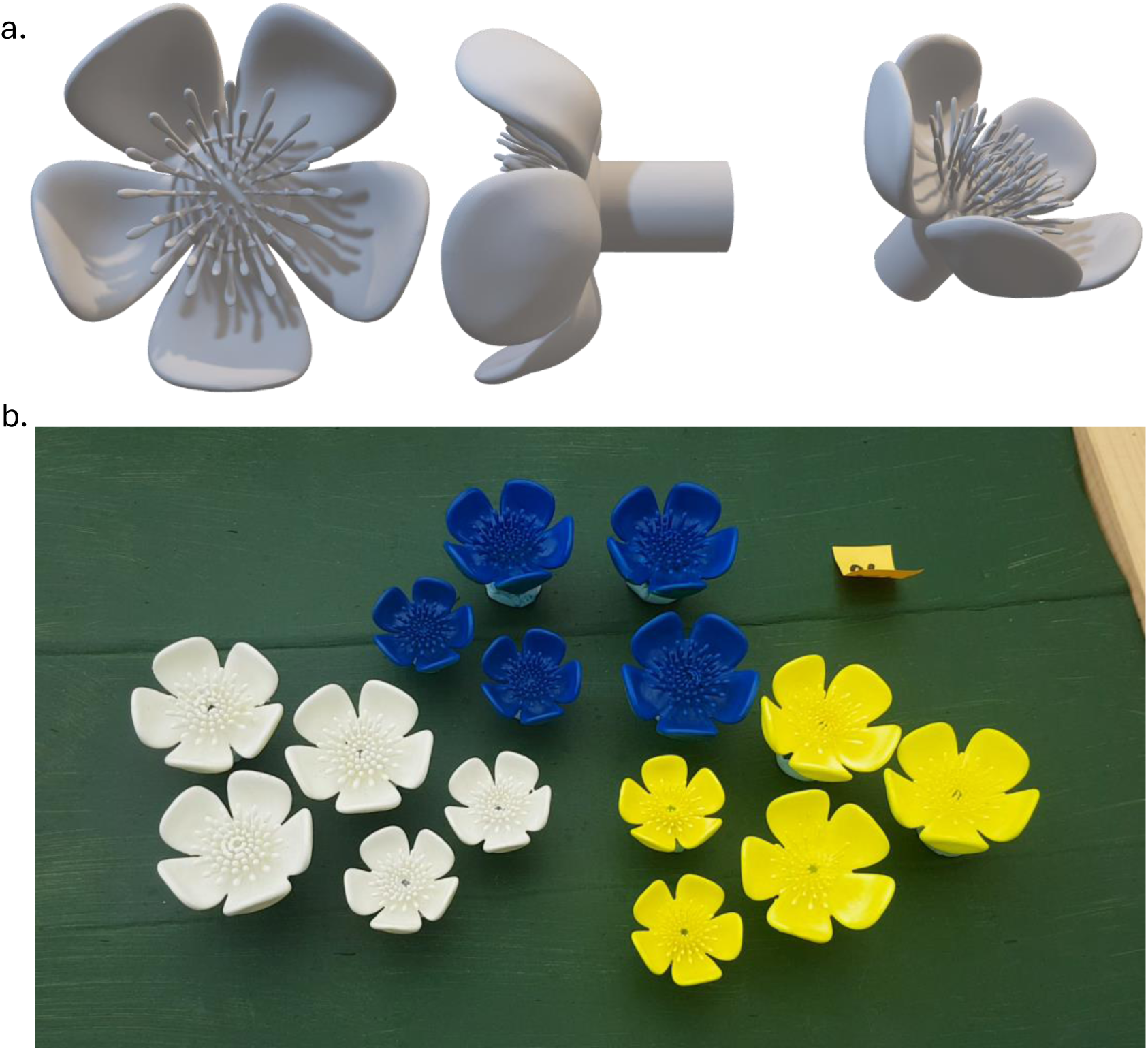
a. Artificial flower model, here showing digital grey colour models from ZBrush. b. Artificial flower set-up in the field. Still taken from one of the filming sessions. The image is closer than that on the pan traps and paper as the video is zoomed in.

The ZBrush file was then converted to stereolithography (STL) format (see Appendix I. Artificial flower STL files), used to represent 3D models in preparation for 3D printing. The STL file was then formatted on ChituBox (Chitu Systems 2024), where the flower size could be adjusted, and multiple flowers could be printed concurrently. After reviewing early prototypes printed at different sizes, 30 mm and 50 mm flower diameters were chosen to reflect the size of buttercups as well as to provide a degree of morphological variety with larger flowers to attract larger insects. Flowers were printed using a resin printer (Brand: Elegoo - Shenzhen, China; Model: Saturn 3) to achieve the most reliable printing of small-scale features such as stamina, as well as to provide a smoother finish. UV fluorescent colours were added to the flowers using *Blacklight UV-Reactive Neon Fluorescent Dayglo Spray Paint* in blue, yellow, and white supplied by UVgear.co.uk. Each flower was first primed using a coating of *Citadel Colour White Scar Priming Spray Paint*, painted using the Dayglo spray paint and sealed using *Clear Varnish top coat spray paint - no UV blocker* by UVgear.co.uk. While all sides of the flowers received some paint due to the aerosol nature of spray paint, the top of the flowers received a more solid application than the sides. Three 50 mm flowers and two 30 mm flowers of each colour were used, leading to a total of five flowers per colour. Flowers were set against a background consisting of a wooden storage box (KNAGGLIG, sourced from IKEA) placed upside down with a rectangular section of cardboard cut to fit the base (28x38cm). The cardboard was painted green using *Wilko Let’s Create green water based acrylic paint*. The flowers were then affixed to the background using Blu Tack in groups corresponding with their colour (Figure 1).

### Pan Traps

Pan traps were constructed from plastic bowls that had approximately the same visual surface area as the other attractants and were the same as those used within the UK Pollinator Monitoring Scheme (UK Pollinator Monitoring Scheme 2025). Each was primed, painted and sealed with the same paints and topcoat used on the artificial flowers. The inside of the pan traps received a more solid application of spray paint than the outside surface. One pan trap of each colour was used to form one replicate, as the average surface area of the five artificial flowers equated to one pan trap (bowl). Unlike traditional surveys in which pan trap bowls are typically filled with water to retain the insects sampled, our pan traps were kept dry to keep the experiment non-lethal (O’Connor et al. 2019). The same methods for the wooden base and green background were used, as for the artificial flowers (Figure 2).

**Figure 2.**
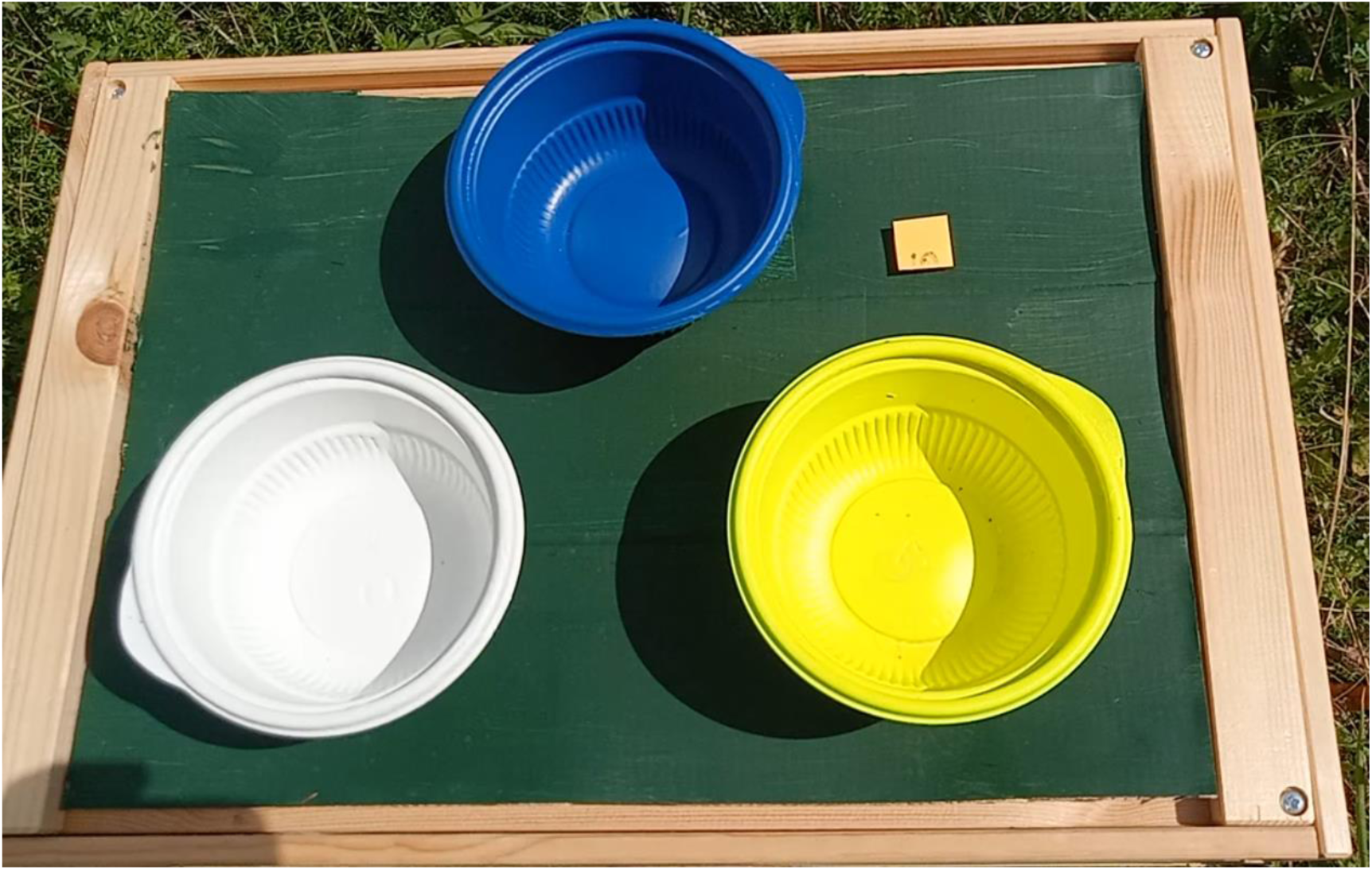
Pan trap attractant set-up. Still taken from one of the filming sessions.

### Coloured Paper

Paper card was cut into 12x12cm squares (being equal in area to the pan trap bowls and artificial flowers), and primed, spray painted and sealed with the same paints and topcoat as used on the pan traps and artificial flowers, to act as the non-lethal equivalent of using sticky traps. The same methods for the wooden base and green background were used, as for the artificial flowers (Figure 3).

**Figure 3.**
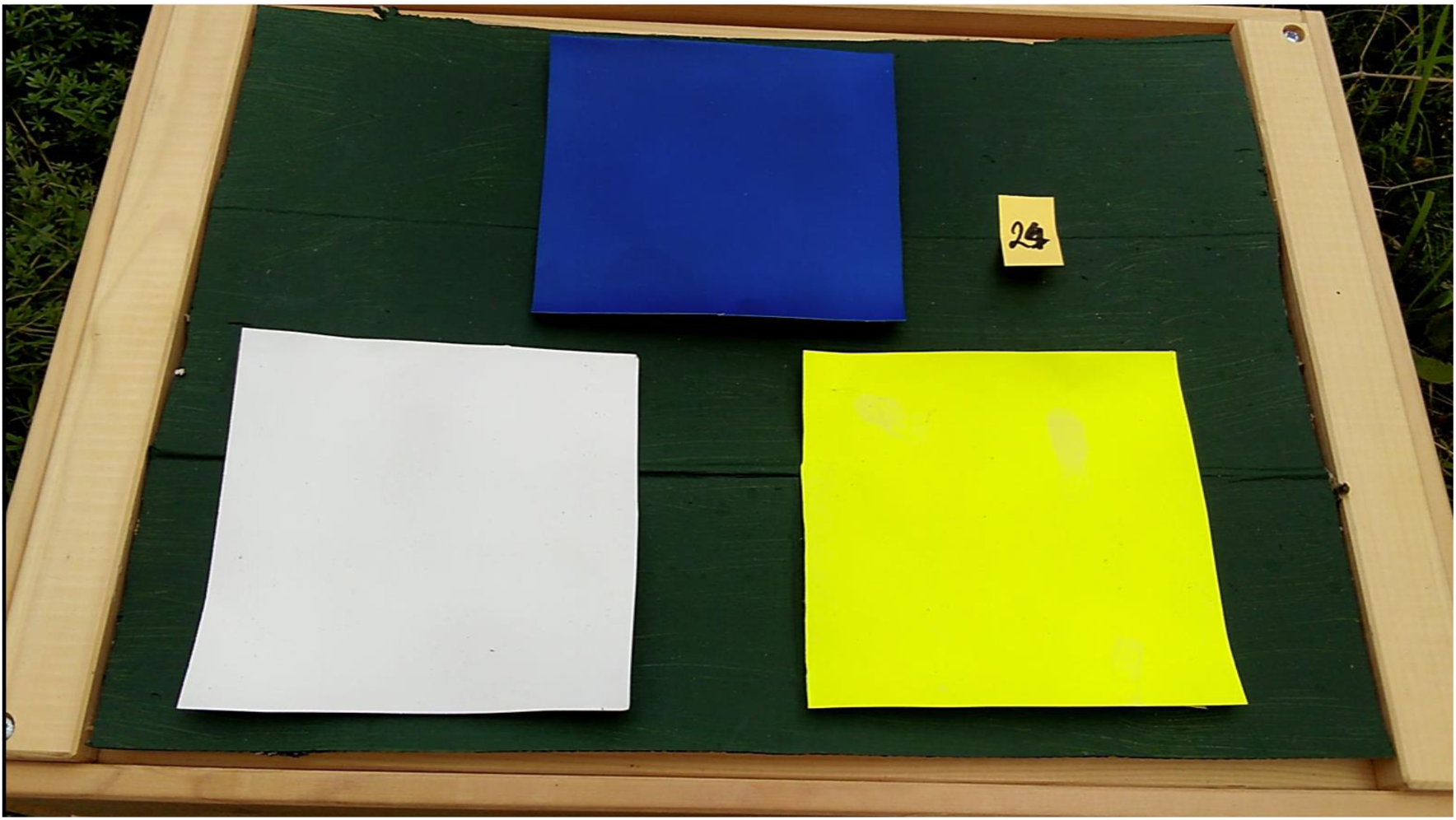
Paper attractant set-up. Still taken from one of the filming sessions.

### Experimental Set-Up

The experiment was undertaken in a well-established wildflower meadow and adjacent grassland on the grounds of the UK Centre for Ecology and Hydrology research station in Oxfordshire, UK (51.602583o N, -1.110813 ° E), between 25^th^ July and 5^th^ September 2023. Filming occurred when the weather was dry, calm and above a minimum temperature of 18 degrees Celsius, which occurred on 17 days in total. The artificial flower, pan trap and coloured paper stations were set out in random spatial configuration relative to each other each day to prevent location bias. Data was collected using three cell phone cameras capturing imagery at a minimum of 720p and 30fps (this experiment used a Nokia TA-1322, a Note 13P Ulefone, and a Samsung Galaxy S7). These devices recorded 57 minutes of data at a time, due to technical constraints of the Note 13P. In total approximately 2-4 hours of data was recorded each day between the hours of 09:00 and 17:00. The variation in time recorded was due to weather and availability. The cell phones were held using tripods with cell phone mounts placed at an appropriate height above each attractant, considered to be sufficient to ensure the attractant could be fully observed in the video with as little of the outer edges of the box showing as possible (Figure 4).

**Figure 4.**
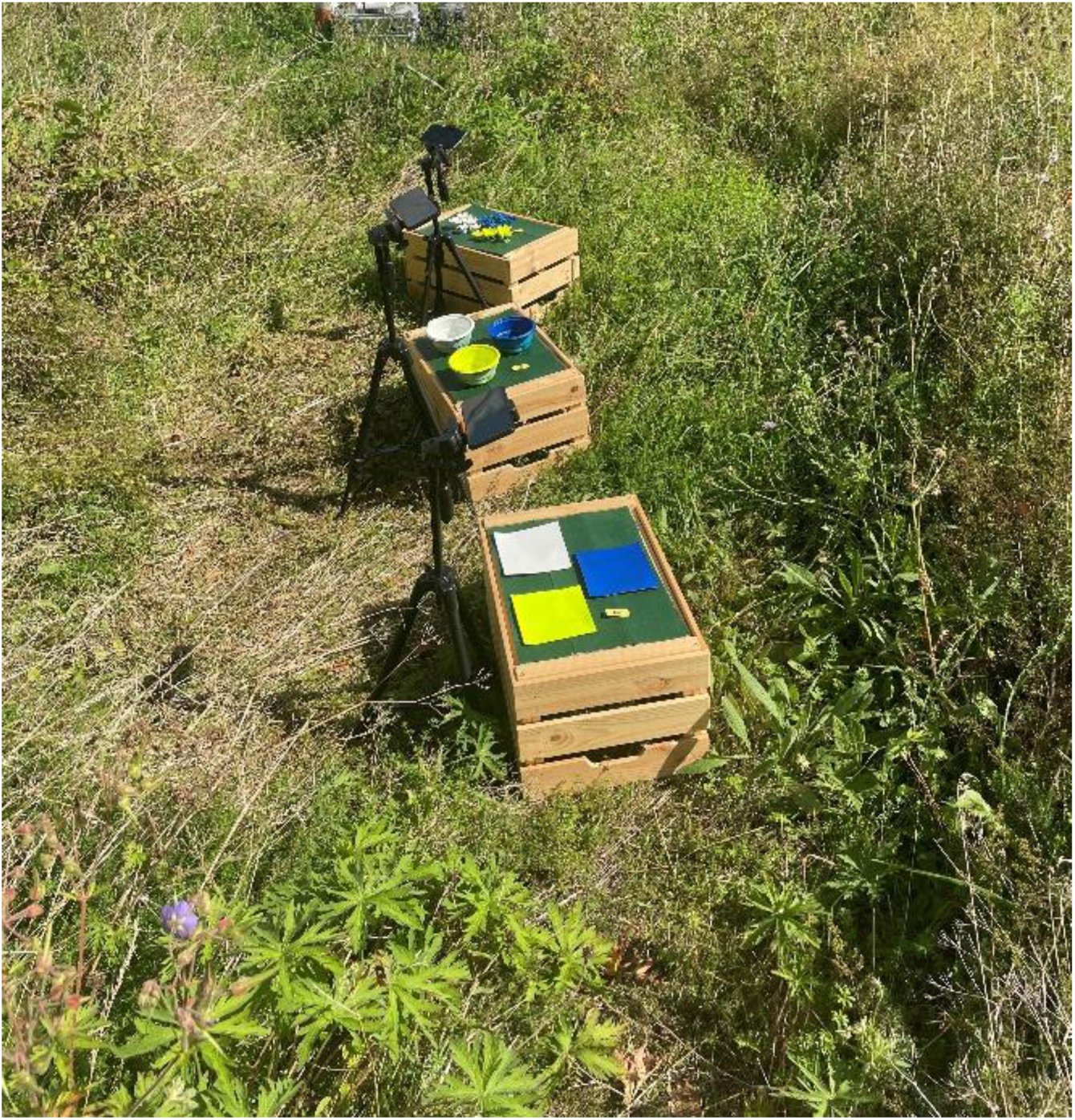
Experimental set up of the attractants (from top to bottom: artificial flowers, pan traps, paper), cell phones, and tripods.

### Data Processing

The three cameras captured a total of 96 hours of video data across all three attractants. All videos were reviewed manually and insect behavioural responses in the videos were recorded and classified as either (i) a landing (coming into physical contact with an attractant, not including landing on the green background) or (ii) an approach only (visibly changing flight behaviour by moving closer to an attractant or slowing down without landing). If an insect landed, its approach was disregarded creating a distinct separation between the landing and approaching behavioural responses. Each insect was identified to species group level following the protocol used by the UK Pollinator Monitoring Scheme FIT Count survey (<u>PoMS FIT Counts)</u> with the groups being: bumblebees, honeybees, solitary bees (all bees other than honeybees and bumblebees), wasps, hoverflies, ‘other flies’ (all flies excluding hoverflies), butterflies and moths, beetles, small insects (insects < 3mm), and ‘other insects’ (insects that did not fall under any other category such as sawflies or true bugs) (UK Pollinator Monitoring Scheme 2024). The attractant, colour, and aggregated duration of the behavioural responses were recorded. Thus, in the case of artificial flowers, where an insect landed on one white flower for 2 seconds, then another white flower for 3 seconds, and then a blue flower for 2 seconds, the total time of the behavioural response recorded would be 5 seconds spent on white and 2 seconds spent on blue. Time spent travelling between flowers was not counted. If an insect left the field of view and returned, it was considered a separate behavioural response.

### Analysis

Data analysis was undertaken using RStudio (v2026.1.0392) (Posit Team 2026). Negative binomial general linear mixed models (GLMMs), using the package glmmTMB, were chosen to analyse the behavioural response results due to the high number of non-responses (0) and single responses (1) per filming session, colour and attractant (Brooks et al. 2025). Overall, the negative binomial model is considered a good-fit for abundance counts where it is expected to have a high percent of zero counts. The time insects spent on the attractants was analysed using a hurdle model (R package: pscl) to separate landings (1) and non-landings (0) using a binomial model and perform a negative binomial regression analysis on the length the insects spent landed on the attractants (Jackman et al. 2024). The model formula included interactions with attractant, insect and colour and the results consider both the likelihood of an insect to land as well as the duration for which it is landed. Further post-hoc testing was done to both the negative binomial GLMM and hurdle model (R package: emmeans) in order to make a multi-way comparison between the different variables and variable interactions (Lenth et al. 2025).

### Comparing Attractants and Colour Preferences

Data was organised by filming session (experiment number), filming session length (experiment length), attractant, behaviour, insect, and colour (see Appendix II. Data Excel file). The interactions with each attractant, colour and behaviour between all filming sessions were compared using the negative binomial GLMMs. The full dataset was analysed with filming session included as random intercept effects and experiment length included as an offset to allow consideration of the overall attractant and colour preferences across the experiment.

The insect groups were then analysed separately to better understand their individual preferences. Insect groups were removed from analysis when the running of a GLMM on that particular group resulted in errors due to the low number of behavioural responses. This resulted in groups with less than 30 behavioural responses being removed from individual analysis. As with the overall analysis, the number of behavioural responses was used as the independent variable and the dependent variables were the attractants, colours, behaviours, and interactions between colour and attractant, behaviour and attractant, and behaviour and colour. The filming session number was included as a random intercept effect to account for weather variations between filming periods and the experiment length was included as an offset.

### Comparing Landing Duration

The time spent landed on the attractants did not include the approach behaviour as the outcome we sought was a determination as to whether insect groups would land and stay longer on specific attractants, colours, or combinations of both. Insect groups were dismissed if there were fewer than ten landings recorded as this led to errors within the hurdle model. To determine significant differences between the time spent landed, the individual length of time an insect spent on one specific attractant and colour for each visit in each experimental session was used as the independent variable. The predictor variables were the attractants, colours, and insects. The interactions were between the attractants and colours, insects and attractants, and colours and insects. The same formula was used for both the binomial model and negative binomial regression.

## Results

### Overall response to attractants and colour

A total of 1725 insect behavioural responses were recorded with 550 approaches and 1225 landings (Figure 5). Artificial flowers received 640 responses, pan traps received 648, and paper received 487 responses. When investigating which attractant and colours the insects preferred overall, our results demonstrated a significant preference for landing on artificial flowers over solid coloured paper (*P = 0.011*) (see Appendix III. Table 2). Furthermore, insects significantly preferred approaching yellow compared to blue (*P < 0.01*) and white (P < 0.001) attractants and significantly preferred landing on white over blue attractants (*P < 0.001*) and on yellow over both blue (P < 0.001) and white (P < 0.001) attractants (see Appendix III. Table 4). Insects landed significantly more on blue flowers over blue pan traps (*P = 0.020*) and paper (*P = 0.017*) and landed significantly more on yellow pan traps (*P = 0.039*) than yellow paper (see Appendix III. Table 1).

**Figure 5.**
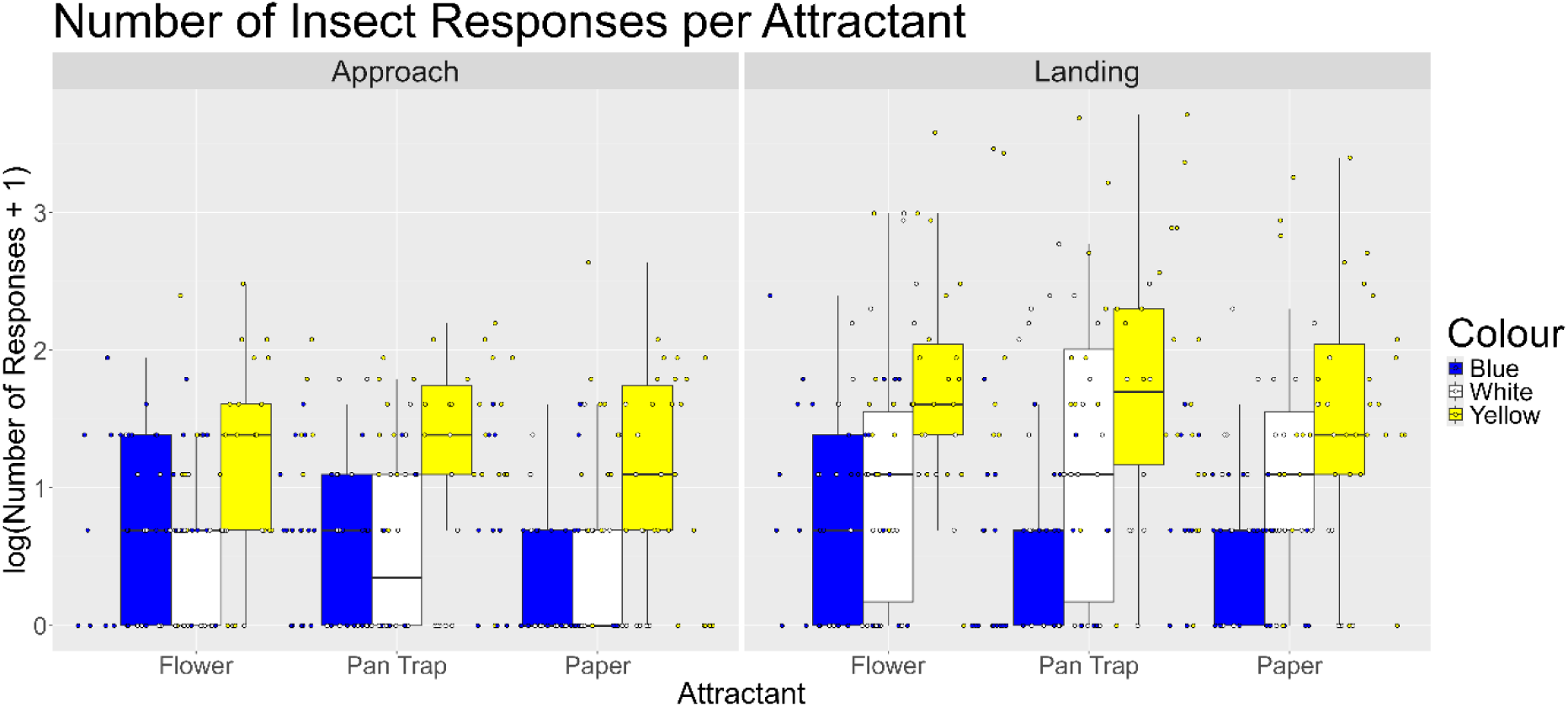
Total insect behavioural responses with each attractant and colour per experiment. Each point represents the total number of times an insect approached or landed on an attractant or colour within one experiment session. The y-axis shows the logged number of behavioural responses to better visualise the results. Each point represents the total number of times an insect responded to the attractant or colour within one experiment session. The box represents the interquartile range of the data, the horizontal black line represents the median, the vertical black lines represent the minimum and maximum values. Any point outside of this range is an outlier.

### Insect-group specific analysis of responses to attractant and colour

As detailed in the PoMS FIT Count protocol, the insects were organised into ten groups. Of these groups, three (honeybees, beetles, butterflies and moths, and other insects) did not have enough responses to analyse using negative binomial GLMMs (Figure 6).

**Figure 6.**
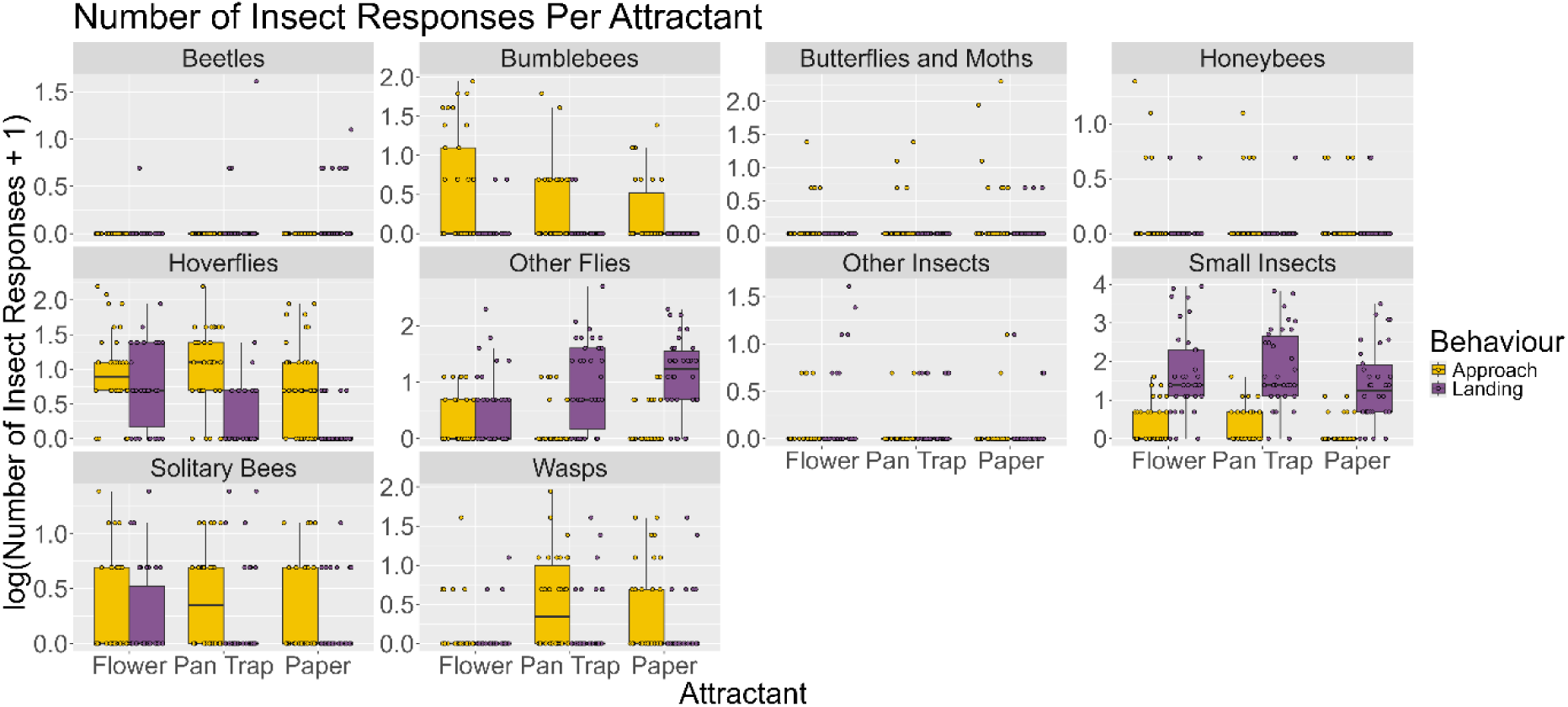
Insect responses per attractant and behaviour (Showing both Landing and Approach only). Each point represents the total number of times an insect approached or landed on an attractant within one experiment session. The y-axis shows the logged number of behavioural responses to better visualise the results. Each point represents the total number of times an insect responded to the attractant or colour within one experiment session. The box represents the interquartile range of the data, the horizontal black line represents the median, the vertical black lines represent the minimum and maximum values. Any point outside of this range is an outlier.

### Attractant Preference

Of the six groups that underwent analysis (see Appendix III. Table 6), five showed a significant preference for specific attractants. Hoverflies landed significantly more on the artificial flowers than the pan traps (*P < 0.001*) and paper (*P < 0.001*). Bumblebees showed a preference for interacting with the artificial flowers compared to paper (*P = 0.043*). Wasps significantly approached pan traps (*P = 0.003*) and paper (*P = 0.025*) more than artificial flowers. Small insects approached artificial flowers significantly more than paper (*P = 0.022*) and landed significantly more on the artificial flowers than pan traps (*P = 0.037*) and paper (*P < 0.001*). ‘Other Flies’ had a significant preference for landing on pan traps (*P = 0.002*) and paper (*P < 0.001*) compared to the artificial flowers.

### Colour Preference

Of the seven groups analysed, five groups showed preference for specific colours (Figure 7, see Appendix III. Table 5 and 7). Hoverflies showed a significant preference for yellow (*P < 0.001*) over white and blue when both approaching and landing. They also exhibited a significant preference for landing on yellow, white, and blue artificial flowers compared to the same-coloured pan traps and paper (Yellow: *P < 0.001* and *P < 0.001*; White: *P = 0.008* and *P < 0.001*; Blue: *P = 0.003* and *P < 0.001*). Bumblebees showed a significant preference for blue over white (*P = 0.01*) as well as yellow over white (*P = 0.049*) with specific preferences for blue artificial flowers over blue paper (*P = 0.018*).

**Figure 7.**
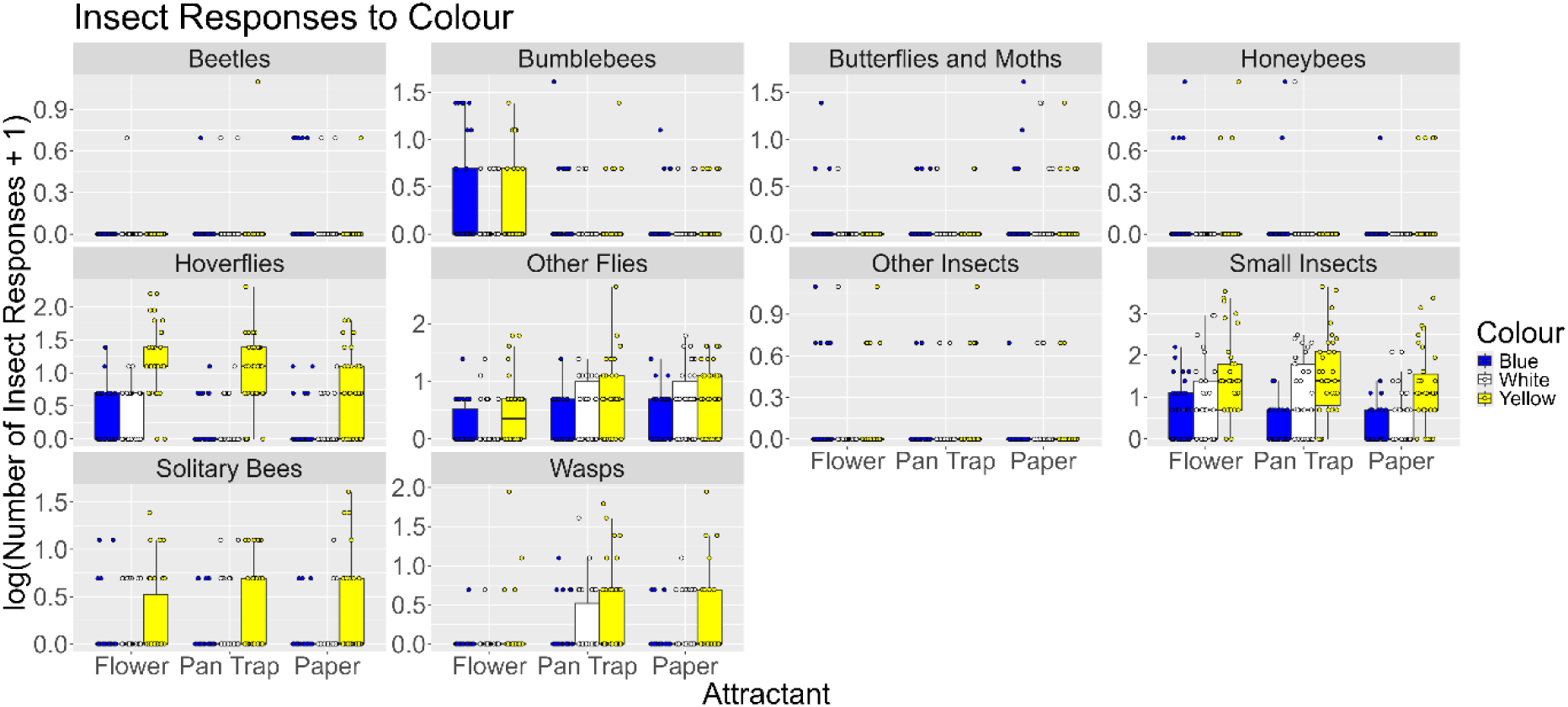
Colour preference of each insect group for each attractant with approach and landing responses combined. The y-axis shows the logged number of responses to represent better visually the different in colour preference. Each point represents the total number of times an insect responded to an attractant or colour within one experiment session. The box represents the interquartile range of the data, the horizontal black line represents the median, the vertical black lines represent the minimum and maximum values. Any point outside of this range is an outlier.

Small insects exhibited a significant overall preference for approaching and landing on yellow compared to blue (Approaching: *P < 0.001;* Landing: *P < 0.001*) and white (Approaching: *P = 0.02;* Landing: *P < 0.001*). They also preferred landing on white attractants over blue ones (*P < 0.001*). Analysis on the interactions between attractants, behaviour and colour showed that small insects significantly preferred approaching and landing on blue artificial flowers over blue pan traps (Approaching: *P = 0.038*; Landing: *P = 0.003*) and paper (Approaching: *P = 0.015*; Landing; *P = 0.006*) as well as approaching and landing on white artificial flowers (Approaching: *P = 0.019*; Landing *P < 0.001*) and pan traps (Approaching: *P = 0.019*; Landing: *P < 0.001*) more than white paper. They also landed more on yellow pan traps than yellow paper (*P = 0.034*).

Solitary bees showed a significant overall preference for approaching and landing on yellow compared to blue (Approaching: *P = 0.026*; Landing: *P = 0.003*) and white (Approaching: *P = 0.024*; Landing; *P = 0.005*) but did not show any preferences for specific colour and attractant combinations. Wasps had a significant preference for approaching and landing on yellow over blue (Approaching: *P = 0.005*; Landing: *P = 0.007*) as well as landing on yellow over white (*P = 0.005*). They have a specific preference for approaching white pan traps over white artificial flowers (*P = 0.041*).

‘Other Flies’ exhibited a significant preference for approaching and landing on yellow) over blue (Approaching: *P = 0.011*; Landing: *P < 0.001*) and white (Approaching: *P = 0.005*; Landing; *P = 0.002*) as well as landing on white over blue (*P = 0.008*). They had specific preferences for landing on yellow and white pan traps and paper over yellow (Pan Traps: *P = 0.002*; Paper: *P = 0.016*) and white (Pan Traps: *P = 0.015*; Paper: *P < 0.001*) artificial flowers.

### Time Spent on each Attractant and Colour

The analysis of time spent on each attractant focused on landing responses to address our hypothesis that insects would spend more time landed on the artificial flowers. Of the ten insect groups, seven groups (beetles, hoverflies, ‘other flies’, ‘other insects’, small insects, solitary bees, and wasps) had enough landing data points for analysing time spent on each attractant and colour and four of the groups spent a significant amount of time on one or more of the attractants and/or colours (Figure 8).

**Figure 8.**
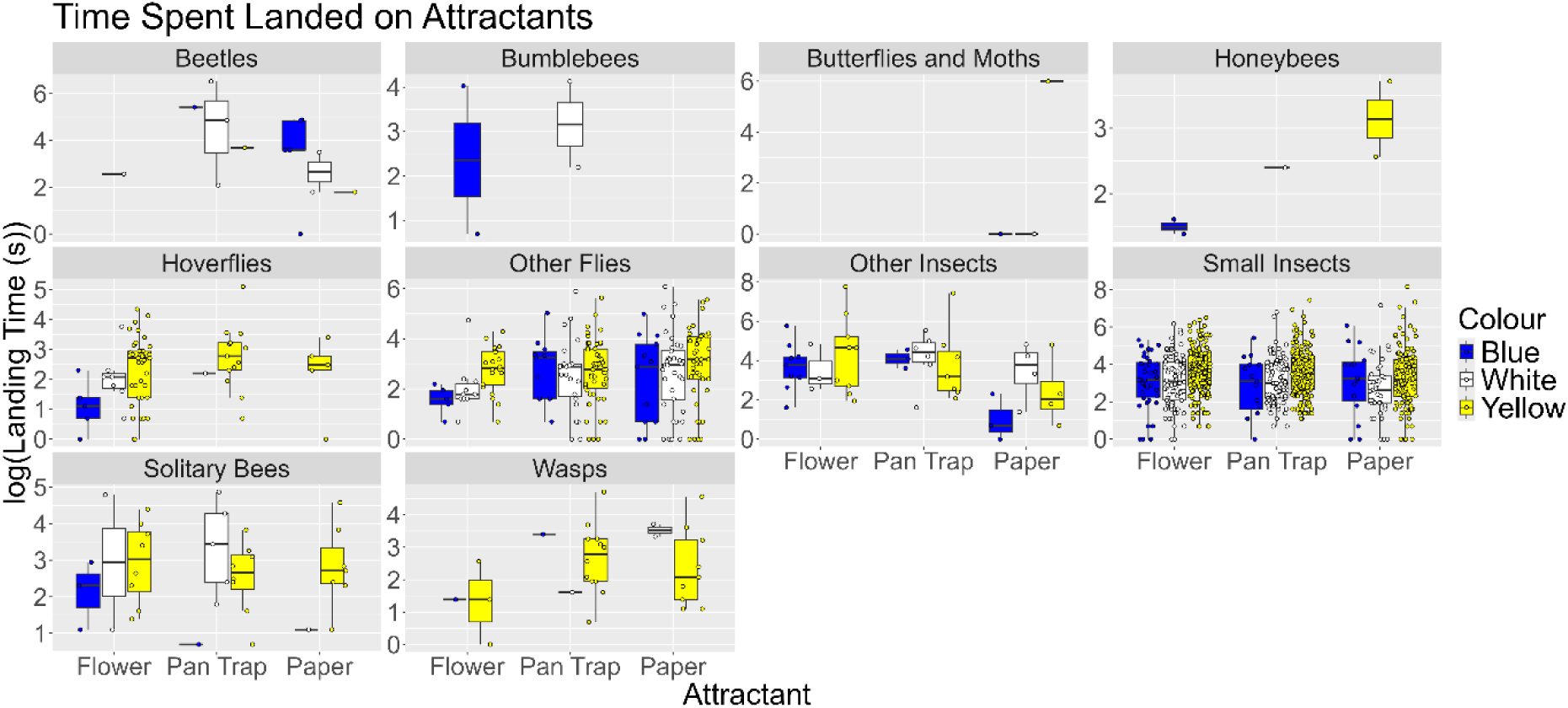
Time each insect spent landed on different attractants and colours. The y-axis shows the length of time spent on the attractants. Each point represents the length of time of an individual insect’s landing. The box represents the interquartile range of the data, the horizontal black line represents the median, the vertical black lines represent the minimum and maximum values. Any point outside of this range is an outlier.

The post-hoc tests (see Appendix III. Table 8, 9, and 10) on the hurdle model showed that hoverflies spent significantly longer landed on the colour yellow than white (*P < 0.001*) or blue (*P < 0.001*). Hoverflies spent significantly longer on the artificial flowers than the paper (*P < 0.001*) but the landing duration did not differ significantly between the artificial flowers and pan traps. When looking at attractant and colour together, hoverflies spent significantly longer on the yellow artificial flowers than the yellow paper (*P < 0.001*).When considering the estimates, despite not spending a significantly different duration on the artificial flowers compared to the pan traps, hoverflies overall do spend more time on average on the artificial flowers than the pan traps (2.178 seconds).

‘Other flies’ spent significantly longer on the yellow (*P < 0.001*) and white (*P = 0.003*) attractants compared to the blue and spent longer overall on the pan traps (*P < 0.001*) and paper (*P < 0.001*) compared to the artificial flowers. For specific attractant and colour interactions, ‘other flies’ spent significantly longer on the yellow and white pan traps (Yellow: *P < 0.001*; White: *P < 0.001*) and paper (Yellow: *P < 0.001*; White: *P < 0.001*) compared to the yellow and white artificial flowers. They also spent significantly longer on the blue paper (P = 0.037) compared to the blue artificial flowers.

Small insects spent significantly longer on the yellow (*P < 0.001*) and white (*P = 0.010*) attractants compared to the blue as well as longer on yellow than white (*P < 0.001*). Solitary bees spent longer landed on the yellow (*P = 0.015*) attractants than the blue. Wasps spent significantly longer on the yellow attractants (*P = 0.017*) compared to the blue. Small insects, solitary bees, and wasps did not exhibit a significant different in landing duration between the attractants, however, the estimates show that solitary bees overall spend at more time on the artificial flowers than the pan traps (1.323 seconds) and paper (2.105 seconds). Further investigation into the estimates of the attractant and colour interactions show that solitary bees spend on average longer on yellow artificial flowers compared to yellow pan traps (2.463 seconds) and paper (3.582 seconds).

## Discussion

We found that key pollinating insect groups showed a preference for artificial flowers over standard attractants, and that they spent longer on artificial flowers when they landed, providing support for the usage of artificial flowers as a candidate attractant with automated monitoring systems.

### Behavioural Observations

Due to the method employing filming of the insect visits, it was possible to observe behavioural differences between the behavioural responses of insects with the different attractants. These observations could help inform future studies using artificial flowers with this method of recording for wild insect behavioural research. The addition of a stamen in the artificial flowers showed insects extending their proboscis (see Appendix IV. Hoverfly landing video) as well as visiting multiple flowers (see Appendix V. Solitary bee landing video). While these observations do not constitute quantifiable data in this experiment, it does open the possibility to explore this further in future studies researching wild insect behaviour using similar methods; for example, comparing the effects of artificial flower size on the behaviour of visiting insects of various sizes.

### Attractant Preferences

We posited a hypothesis that insects would prefer interacting with artificial flowers than with other more ‘traditional’ attractants used in pollinator monitoring. Five out of seven insect groups analysed exhibited a strong preference for either specific attractants or colours (hoverflies, bumblebees, small insects, and ‘other flies’). Of these five groups, three (hoverflies, bumblebees, and small insects) showed a preference for artificial flowers, and two (‘other flies’ and wasps) showed a preference for pan traps and paper. Bumblebees, with only five landings across all the attractants, did approach the artificial flowers significantly more than the paper. The small insects consistently overall approached and landed more on the artificial flowers. The hoverflies had very a similar number of approaches across the attractants but a high number of landings on artificial flowers showing an increase in abundance. This result is key as it shows that in addition to the colour yellow attracting hoverflies, the artificial flower shape led to more interest in landing than the pan traps or paper. This landing behaviour is necessary for the insects to be easily identified under the insect camera traps as often the insect needs to be relatively still or slow (up to 2 seconds) to be fully detected by certain cameras such as InsectDetect (Sittinger et al. 2024).

With the exception of the ‘other flies’ group and wasps, artificial flowers were not found to be significantly less effective than the other two attractants. Therefore, the use of artificial flowers as attractants under the insect camera traps, given the evidence of their superior ability to attract bumblebees, hoverflies, and small insects, was not found to negatively impact observations of other groups. When comparing how the attractants appeared under the camera, the artificial flowers and paper had the benefit of not producing much shadow on the top face of the attractant most visible to the camera while the pan traps often cast shadows inside the bowl, disrupting the viewers’ or camera’s ability to see the whole bowl clearly. When considering this, and with the knowledge that artificial flowers attracted more insects overall than the paper, artificial flowers show significant potential as an attractant for automated monitoring systems.

### Colour Preferences

We hypothesised that the insect groups would exhibit colour preferences and would be more likely to interact with the colours that align with preferences described in behavioural research. Based on the current literature on colour preferences in pollinating insects, hoverflies of the genus *Eristalis* exhibit an innate preference for yellow and *Bombus terrestris* bumblebees have an innate preference for blue/UV blue (Raine & Chittka 2007; An et al. 2018). Small insects and ‘other flies’ fall under the generalist category with the generalist colours being considered yellow and white (Lunau 2014). All these established preferences were supported with significant results in the present study (see Appendix III. Table 7). While innate colour preferences are important to note, we cannot ignore the learned colour preferences from the floral selection in the experiment’s surrounding area. Overall, this indicates that using the correct colours to attract specific insect groups is key when designing artificial flowers.

### Time Spent on each Attractant and Colour

The longer an insect stays on an attractant the more likely it is to be registered by a camera trap either through AI detection or timelapse. Therefore, while attraction is important, so too is an attractant’s ability to retain an insect on its surface. Both methods of data collection using an insect camera trap, AI and timelapse, need the insect to stay on the background for a few seconds to ensure the camera can take an adequate image. For timelapse, this ensures the insect is indeed captured by the camera whilst in an AI-based detection model, it ensures the insect stays in one place long enough to get a clear image.

Alongside the knowledge that hoverflies significantly preferred landing on the artificial flowers compared to the pan traps and paper, it is important to note that they also spent on average longer on the artificial flowers than the pan traps and paper even though only the difference between artificial flowers and paper was significant.

This means that using the artificial flowers for automated monitoring will increase the number of hoverflies that land as well as how long they stay landed ensuring that the camera traps will detect them. Additionally, the solitary bees, whilst showing no significant difference in preference between attractants, spent longer on the yellow and spent 2-3 more seconds on average on the yellow artificial flowers than the yellow pan traps and paper, so could be easily detected by the camera traps if using the artificial flowers. This is crucial, especially considering that insects are more likely to be detected by cameras such as the InsectDetect if they are under the camera for more than 2 seconds (Sittinger et al. 2024). Therefore, using the artificial flowers, specifically the yellow ones, could ensure higher probability of detection of key pollinators, hoverflies and solitary bees.

### Wider Context

Two groups that showed strong preferences for the artificial flowers (hoverflies and bumblebees) are among some of the more common pollinating groups known to have significant impact on pollination (Doyle et al. 2020; Khalifa et al. 2021). Hoverflies are key pollinators that visit up to 72% of global food crops and are worth approximately $300 billion in pollination services (Doyle et al. 2020). They provide further ecosystem services such as organic waste recycling and crop pest protection (Doyle et al. 2020). Given the results presented, increased landings observed, and greater time spent on the artificial flowers, the use of the artificial flowers and insect camera traps indicates that this approach could be a viable, non-lethal, method for monitoring this key group.

Bumblebee populations have been declining globally as noted by an overall decrease of 22.5% in 2024 of British bumblebees recorded by the Bumblebee Conservation Trust (Comont & Dickinson 2024, 2025), projected declines of European bumblebees (Ghisbain et al. 2024), and a decline of the Western bumblebee (*Bombus occidentalis*) in the United States (Graves et al. 2020). This is concerning as bumblebees play a pivotal role in wild pollination and are commonly used in commercial pollination (Khalifa et al. 2021). With the decline in bumblebees, it is important to maintain high levels of population monitoring, which, alongside pollinator monitoring schemes, could be undertaken using artificial flowers and insect camera traps. Spatial gaps in data samples for bumblebee monitoring (such as for the Western Bumblebee) shows how automated monitoring using artificial flowers could offer a useful tool in creating wider spatio-temporal data gathering methods (Graves et al. 2020). Whilst the bumblebee behavioural response with the artificial flowers primarily comprised only approaches, our method provides a useful foundation to begin including additional attraction cues that could ensure a higher rate of conversion, from approaches to landings.

The EU Nature Restoration Law 2024 describes how monitoring pollinator diversity and populations is an action that will help reverse the decline of pollinator populations (European Union 2025). Already, projects such as ORBIT (Taxonomic Resources for European Bees), and TAXO-FLY (Taxonomic Resources for European Hoverflies) have been developed to help generate taxonomic information for improved pollinator monitoring (Potts et al. 2024). Knowing that artificial flowers can increase visits to automated pollinator monitoring systems can thus benefit directly the goals set out by the EU Nature Restoration Law 2024.

Furthermore, when considering the use of artificial flowers with AI-assisted, image-based monitoring methods, it’s worthwhile to consider the benefits and potential drawbacks of such a system. The benefits of using artificial flowers in AI-assisted monitoring include being able to have a fully controlled system that produces no noise introduced by looking at natural flowers. Additionally, artificial flowers require little to no maintenance compared to live flowers that experience senescence and move day to day, and using the artificial flower models can ensure monitoring efforts are consistent across sites and regions by using a standardised attractant. With the freedom of 3D modelling, it is possible to customise artificial flowers to attract species of interest in a way that is not possible with pan traps and coloured paper. The drawbacks, however, are that artificial flowers are very likely not as attractive as real flowers and insect responses to them do not represent true ‘pollination’. Finally, responses to the artificial flowers could be biased by the learned preference of local insects. This is why it is important to continue researching the use of artificial flowers for AI-assisted, imaged based monitoring methods as there are many more avenues to explore before artificial flowers can be a standardised attractant.

### Next Steps

The present study was limited to observation in the late-summer period in northern Europe. While we did find general patterns within species groups that shows that using artificial flowers could be useful in monitoring certain groups (e.g. hoverflies and bees), we suggest the validation of these findings be tested in new geographic regions or time periods.

Here we presented one, relatively simple, 3D model of a flower. Knowing that a complex floral shape as well as other cues (e.g. scent) can influence increased behavioural response in some insect groups naturally invites the exploration of more complex artificial flowers for future automated monitoring (Wilmsen et al. 2017; Nordström et al. 2017; Chapman et al. 2023). As some insect groups in this study approached the artificial flowers much more than they landed, we suggest future studies explore the inclusion of other attraction cues such as scent and nectar guides (both human visible and UV) (Leonard & Papaj 2011; Knauer & Schiestl 2015; Wilson et al. 2016; Hirota et al. 2019), as well as less studied cues such as floral temperature, humidity, and electrostatic fields (Clarke et al. 2013; Harrap et al. 2017). This could be useful in improving passive monitoring for bumblebees or butterflies which might respond more to the artificial flowers with added attraction cues.

Future studies should also explore designing artificial flowers for specific insect groups (e.g. bumblebees and butterflies) based on the established attraction cue preferences in the literature for these specific groups such as flower morphology (Krishna & Keasar 2021), colour, UV fluorescence (Papiorek et al. 2016), and scent (Reddi & Bai 1984; Chapman et al. 2023). These taxa specific designs could be used to monitor specific insect groups of interest or groups that are more rare, as well as creating a more robust method of passively monitoring the insects not well represented by other methods (Baum & Wallen 2011; Vrdoljak & Samways 2012; Hudson et al. 2020). As insect diversity was not possible to fully determine due to the nature of using broad groups, future studies could explore this. Additionally, further studies could focus on the benefits and drawbacks of using different types of artificial flowers under AI assisted image-based monitoring systems rather than recorded by video, where it will be critical to develop an understanding of the taxonomic resolution that can be achieved.

They could also explore the effects of shadows made by the artificial flowers, the number of artificial flowers used, and how the orientation of the insect landing on the artificial flower impacts the insect detection.

## Conclusions

In this experiment, we show that artificial flowers can serve as efficient and consistent attractants for automated monitoring, compared with current attraction methods. The use of artificial flowers with automated monitoring systems could lead to more widespread standardised non-lethal monitoring of key pollinating insect populations in various ecosystems. Covering a wider spatio-temporal area, these systems could ensure earlier interventions in pollinating insect conservation efforts, informing policy development and allowing these insects to maintain necessary ecosystem services.

## Supporting information

Appendix 1. Artificial flower STL file

Appendix 3. Results tables 1-10

Appendix 4. Video of a solitary bee landing on the artificial flowers

Appendix 5. Video of a hoverfly landing on the artificial flowers

## Appendices

Additional supporting information may be found in the online version of this article:

Appendix I. Artificial flower STL file.
Appendix II. Data Excel file.
Appendix III. Results Tables (1-10)
Appendix IV. Video of a hoverfly landing on the artificial flowers and exploring the stamen with its proboscis.
Appendix V. Video of a solitary bee landing and exploring several artificial flowers.

## Author contributions

A. Ash: Conceptualization, Methodology, Analysis, Investigation, Writing-Original Draft, Visualisation, Validation, Data Curation. S. Hallett, T. August, C. Carvell, L. Williams: Resources acquisition, Project Administration, Methodology, Writing-Review and Editing, Supervision, Funding Acquisition.

## Acknowledgements

This work was supported by the UK Natural Environment Research Council through the CENTA Doctoral Training Partnership [NERC Ref: NE/S007350/1].

## Data accessibility statement

Raw data supporting this study is available upon request.

## Declaration of competing interest

The authors declare that they have no known competing financial interests or personal relationships that could have appeared to influence the work reported in this paper.

## Notes

### Competing Interest Statement

The authors have declared no competing interest.

