## Appendix 3. Results tables 1-10 for "Testing the efficacy of artificial flowers as a novel attractant for automated pollinator monitoring"

***List of Tables***

**Table 1 – All insects combined analysis of attractant preference when considering interactions with behaviour and colour (~attractant|behaviour|colour)**

**Table 2 – All insects combined attractant preference when considering interactions with only behaviour (~attractant|behaviour)**

**Table 3 – All insects combined analysis of attractant preference when considering interactions with only colour (~attractant|colour)**

**Table 4 - All insects combined analysis of colour preference when considering interactions with only behaviour (~colour|behaviour)**

**Table 5 – Individual insect group analysis of attractant preference when considering interactions with behaviour and colour (~attractant|behaviour|colour)**

**Table 6 – Individual insect group analysis of attractant preference when considering interactions with only behaviour (~attractant|behaviour)**

**Table 7 – Individual insect group analysis of colour preference when considering interactions with behaviour (~colour|behaviour)**

**Table 8 – Hurdle model analysis of time spent landed on the attractants. Analysis of attractant spent longer on when considering interactions with insect groups and colour (~attractant|insect|colour)**

**Table 9 – Hurdle model analysis of time spent landed on the attractants. Analysis of attractant spent longer on when considering interactions with insect group(~attractant|insect)**

**Table 10– Hurdle model analysis of time spent landed on the attractants. Analysis of colour spent longer on when considering interactions with insect group (~colour|insect)**

Table 1. **All insects combined analysis of attractant preference when considering interactions with behaviour and colour (~attractant|behaviour|colour)**

Results from pairwise test on emmeans post-hoc tests on negative binomial GLMMs on all insect groups combined, contrasting the different attractants when considering their interactions with behaviour and colour.

Values in bold signify that there is a significant behavioural response occurring.

P < 0.001 = high significance, P < 0.01 = medium significance, P < 0.05 = low significance.

| *Attractant Contrast* | *Behaviour* | *Colour* | *Estimate* | *SE* | *Z ratio* | *P value* |
| --- | --- | --- | --- | --- | --- | --- |
| Flower - Pan Trap | Approach | Blue | 0.461 | 0.205 | 2.245 | 0.063 |
| Flower - Paper | Approach | Blue | 0.490 | 0.208 | 2.358 | **0.048** |
| Pan Trap - Paper | Approach | Blue | 0.029 | 0.225 | 0.129 | 0.990 |
| Flower - Pan Trap | Approach | White | -0.245 | 0.195 | -1.256 | 0.419 |
| Flower - Paper | Approach | White | 0.068 | 0.208 | 0.327 | 0.942 |
| Pan Trap - Paper | Approach | White | 0.313 | 0.203 | 1.538 | 0.272 |
| Flower - Pan Trap | Approach | Yellow | -0.139 | 0.144 | -0.960 | 0.601 |
| Flower - Paper | Approach | Yellow | 0.154 | 0.156 | 0.988 | 0.583 |
| Pan Trap - Paper | Approach | Yellow | 0.293 | 0.152 | 1.919 | 0.133 |
| Flower - Pan Trap | Landing | Blue | 0.546 | 0.203 | 2.680 | **0.020** |
| Flower - Paper | Landing | Blue | 0.566 | 0.206 | 2.736 | **0.017** |
| Pan Trap - Paper | Landing | Blue | 0.019 | 0.223 | 0.086 | 0.995 |
| Flower - Pan Trap | Landing | White | -0.160 | 0.161 | -0.990 | 0.582 |
| Flower - Paper | Landing | White | 0.143 | 0.171 | 0.837 | 0.679 |
| Pan Trap - Paper | Landing | White | 0.303 | 0.165 | 1.834 | 0.158 |
| Flower - Pan Trap | Landing | Yellow | -0.054 | 0.109 | -0.493 | 0.874 |
| Flower - Paper | Landing | Yellow | 0.229 | 0.116 | 1.966 | 0.120 |
| Pan Trap - Paper | Landing | Yellow | 0.283 | 0.116 | 2.438 | **0.039** |

Table 2. ***All insects combined attractant preference when considering interactions with only behaviour (~attractant|behaviour)***

Results from pairwise test on emmeans post-hoc tests on negative binomial GLMMs on all insect groups combined, contrasting the different attractants when considering their interactions with behaviour.

Values in bold signify that there is a significant behavioural response occurring.

P < 0.001 = high significance, P < 0.01 = medium significance, P < 0.05 = low significance.

| *Attractant Contrast* | *Behaviour* | *Estimate* | *SE* | *Z ratio* | *P value* |
| --- | --- | --- | --- | --- | --- |
| Flower - Pan Trap | Approach | 0.025 | 0.134 | 0.191 | 0.980 |
| Flower - Paper | Approach | 0.237 | 0.142 | 1.666 | 0.218 |
| Pan Trap - Paper | Approach | 0.211 | 0.144 | 1.471 | 0.304 |
| Flower - Pan Trap | Landing | 0.110 | 0.105 | 1.054 | 0.542 |
| Flower - Paper | Landing | 0.312 | 0.109 | 2.862 | **0.011** |
| Pan Trap - Paper | Landing | 0.202 | 0.111 | 1.809 | 0.166 |

Table 3. **All insects combined analysis of attractant preference when considering interactions with only colour (~attractant|colour)**

Results from pairwise test on emmeans post-hoc tests on negative binomial GLMMs on all insect groups combined, contrasting the different attractants when considering their interactions with colour.

Values in bold signify that there is a significant behavioural response occurring.

P < 0.001 = high significance, P < 0.01 = medium significance, P < 0.05 = low significance.

| *Attractant Contrast* | *Colour* | *Estimate* | *SE* | *Z ratio* | *P value* |
| --- | --- | --- | --- | --- | --- |
| Flower - Pan Trap | Blue | 0.510 | 0.187 | 2.725 | **0.017** |
| Flower - Paper | Blue | 0.516 | 0.186 | 2.762 | **0.015** |
| Pan Trap - Paper | Blue | 0.005 | 0.205 | 0.026 | 0.999 |
| Flower - Pan Trap | White | -0.219 | 0.159 | -1.379 | 0.351 |
| Flower - Paper | White | 0.096 | 0.169 | 0.569 | 0.836 |
| Pan Trap - Paper | White | 0.316 | 0.163 | 1.933 | 0.129 |
| Flower - Pan Trap | Yellow | -0.092 | 0.099 | -0.922 | 0.625 |
| Flower - Paper | Yellow | 0.198 | 0.107 | 1.846 | 0.154 |
| Pan Trap - Paper | Yellow | 0.290 | 0.105 | 2.755 | 0.016 |

Table 4. **All insects combined analysis of colour preference when considering interactions with only behaviour (~colour|behaviour)**

Results from a pairwise test on emmeans post-hoc tests on negative binomial GLMMs on all insect groups combined, contrasting the different colour preferences when considering their interactions with behaviour. Each insect group was analysed separately.

Values in bold signify that there is a significant behavioural response occurring.

P < 0.001 = high significance, P < 0.01 = medium significance, P < 0.05 = low significance.

| *Colour Contrast* | *Behaviour* | *Estimate* | *SE* | *Z ratio* | *P value* |
| --- | --- | --- | --- | --- | --- |
| Blue - White | Approach | 0.272 | 0.17 | 1.598 | 0.246 |
| Blue - Yellow | Approach | -0.971 | 0.134 | -7.229 | **P < 0.001** |
| White - Yellow | Approach | -1.244 | 0.146 | -8.468 | **P < 0.001** |
| Blue - White | Landing | -0.853 | 0.134 | -6.345 | **P < 0.001** |
| Blue - Yellow | Landing | -1.717 | 0.122 | -14.014 | **P < 0.001** |
| White - Yellow | Landing | -0.864 | 0.09 | -9.529 | **P < 0.001** |

Table 5. **Individual insect group analysis of attractant preference when considering interactions with behaviour and colour (~attractant|behaviour|colour)**

Results from pairwise test on emmeans post-hoc tests on negative binomial GLMMs of insect count data, contrasting the different attractants when considering their interactions with behaviour and colour. Each insect group was analysed separately.

Values in bold signify that there is a significant behavioural response occurring.

P < 0.001 = high significance, P < 0.01 = medium significance, P < 0.05 = low significance.

| **Insect** | **Contrast** | **Behaviour** | **Colour** | **Estimate** | **SE** | **Z ratio** | **P value** |
| --- | --- | --- | --- | --- | --- | --- | --- |
| Hoverflies | Flower - Pan Trap | Approach | Blue | 0.409 | 0.449 | 0.911 | 0.633 |
|  | Flower - Paper | Approach | Blue | 0.496 | 0.473 | 1.048 | 0.546 |
|  | Pan Trap - Paper | Approach | Blue | 0.086 | 0.519 | 0.167 | 0.984 |
|  | Flower - Pan Trap | Approach | White | 0.210 | 0.462 | 0.455 | 0.891 |
|  | Flower - Paper | Approach | White | 0.396 | 0.513 | 0.771 | 0.720 |
|  | Pan Trap - Paper | Approach | White | 0.185 | 0.545 | 0.340 | 0.938 |
|  | Flower - Pan Trap | Approach | Yellow | -0.144 | 0.183 | -0.786 | 0.711 |
|  | Flower - Paper | Approach | Yellow | 0.165 | 0.199 | 0.826 | 0.686 |
|  | Pan Trap - Paper | Approach | Yellow | 0.309 | 0.194 | 1.590 | 0.249 |
|  | Flower - Pan Trap | Landing | Blue | 1.748 | 0.541 | 3.229 | **0.003** |
|  | Flower - Paper | Landing | Blue | 2.580 | 0.666 | 3.874 | ***P < 0.001*** |
|  | Pan Trap - Paper | Landing | Blue | 0.832 | 0.734 | 1.134 | 0.492 |
|  | Flower - Pan Trap | Landing | White | 1.549 | 0.522 | 2.965 | **0.008** |
|  | Flower - Paper | Landing | White | 2.481 | 0.666 | 3.722 | ***P < 0.001*** |
|  | Pan Trap - Paper | Landing | White | 0.931 | 0.726 | 1.283 | 0.404 |
|  | Flower - Pan Trap | Landing | Yellow | 1.194 | 0.309 | 3.859 | ***P < 0.001*** |
|  | Flower - Paper | Landing | Yellow | 2.250 | 0.475 | 4.734 | ***P < 0.001*** |
|  | Pan Trap - Paper | Landing | Yellow | 1.055 | 0.526 | 2.005 | 0.110 |
| Bumblebees | Flower - Pan Trap | Approach | Blue | 0.924 | 0.422 | 2.186 | 0.073 |
|  | Flower - Paper | Approach | Blue | 1.343 | 0.496 | 2.707 | **0.018** |
|  | Pan Trap - Paper | Approach | Blue | 0.418 | 0.567 | 0.738 | 0.740 |
|  | Flower - Pan Trap | Approach | White | 0.729 | 0.801 | 0.910 | 0.633 |
|  | Flower - Paper | Approach | White | 0.012 | 0.696 | 0.017 | 0.999 |
|  | Pan Trap - Paper | Approach | White | -0.717 | 0.842 | -0.852 | 0.670 |
|  | Flower - Pan Trap | Approach | Yellow | 0.705 | 0.472 | 1.491 | 0.294 |
|  | Flower - Paper | Approach | Yellow | 1.068 | 0.526 | 2.027 | 0.105 |
|  | Pan Trap - Paper | Approach | Yellow | 0.363 | 0.593 | 0.612 | 0.813 |
|  | Flower - Pan Trap | Landing | Blue | 0.097 | 1.092 | 0.089 | 0.995 |
|  | Flower - Paper | Landing | Blue | 22.67 | 38811.530 | 0.001 | 1.000 |
|  | Pan Trap - Paper | Landing | Blue | 22.572 | 38811.530 | 0.001 | 1.000 |
|  | Flower - Pan Trap | Landing | White | -0.097 | 1.092 | -0.089 | 0.995 |
|  | Flower - Paper | Landing | White | 21.338 | 38811.530 | 0.001 | 1.000 |
|  | Pan Trap - Paper | Landing | White | 21.435 | 38811.530 | 0.001 | 1.000 |
|  | Flower - Pan Trap | Landing | Yellow | -0.121 | 1.202 | -0.101 | 0.994 |
|  | Flower - Paper | Landing | Yellow | 22.394 | 38811.530 | 0.001 | 1.000 |
|  | Pan Trap - Paper | Landing | Yellow | 22.516 | 38811.530 | 0.001 | 1.000 |
| Solitary Bees | Flower - Pan Trap | Approach | Blue | -0.006 | 0.594 | -0.01 | 0.999 |
|  | Flower - Paper | Approach | Blue | 0.62 | 0.721 | 0.859 | 0.665 |
|  | Pan Trap - Paper | Approach | Blue | 0.626 | 0.715 | 0.875 | 0.655 |
|  | Flower - Pan Trap | Approach | White | -0.408 | 0.557 | -0.733 | 0.743 |
|  | Flower - Paper | Approach | White | 0.397 | 0.728 | 0.545 | 0.848 |
|  | Pan Trap - Paper | Approach | White | 0.806 | 0.687 | 1.172 | 0.469 |
|  | Flower - Pan Trap | Approach | Yellow | -0.574 | 0.438 | -1.31 | 0.389 |
|  | Flower - Paper | Approach | Yellow | -0.56 | 0.453 | -1.236 | 0.431 |
|  | Pan Trap - Paper | Approach | Yellow | 0.013 | 0.395 | 0.033 | 0.999 |
|  | Flower - Pan Trap | Landing | Blue | 0.355 | 0.712 | 0.498 | 0.871 |
|  | Flower - Paper | Landing | Blue | 1.324 | 0.870 | 1.521 | 0.280 |
|  | Pan Trap - Paper | Landing | Blue | 0.968 | 0.846 | 1.145 | 0.486 |
|  | Flower - Pan Trap | Landing | White | -0.047 | 0.587 | -0.080 | 0.996 |
|  | Flower - Paper | Landing | White | 1.101 | 0.768 | 1.432 | 0.324 |
|  | Pan Trap - Paper | Landing | White | 1.148 | 0.748 | 1.535 | 0.274 |
|  | Flower - Pan Trap | Landing | Yellow | -0.212 | 0.446 | -0.475 | 0.882 |
|  | Flower - Paper | Landing | Yellow | 0.143 | 0.501 | 0.285 | 0.956 |
|  | Pan Trap - Paper | Landing | Yellow | 0.355 | 0.478 | 0.743 | 0.737 |
| Wasps | Flower - Pan Trap | Approach | Blue | -1.776 | 1.097 | -1.618 | 0.237 |
|  | Flower - Paper | Approach | Blue | -1.390 | 1.129 | -1.230 | 0.434 |
|  | Pan Trap - Paper | Approach | Blue | 0.385 | 0.664 | 0.581 | 0.830 |
|  | Flower - Pan Trap | Approach | White | -2.531 | 1.048 | -2.415 | **0.041** |
|  | Flower - Paper | Approach | White | -2.15 | 1.065 | -2.018 | 0.107 |
|  | Pan Trap - Paper | Approach | White | 0.380 | 0.459 | 0.827 | 0.685 |
|  | Flower - Pan Trap | Approach | Yellow | -1.152 | 0.535 | -2.154 | 0.079 |
|  | Flower - Paper | Approach | Yellow | -0.900 | 0.55 | -1.635 | 0.230 |
|  | Pan Trap - Paper | Approach | Yellow | 0.252 | 0.373 | 0.674 | 0.778 |
|  | Flower - Pan Trap | Landing | Blue | -1.631 | 1.267 | -1.287 | 0.402 |
|  | Flower - Paper | Landing | Blue | -1.369 | 1.294 | -1.057 | 0.540 |
|  | Pan Trap - Paper | Landing | Blue | 0.262 | 0.798 | 0.328 | 0.942 |
|  | Flower - Pan Trap | Landing | White | -2.386 | 1.251 | -1.906 | 0.136 |
|  | Flower - Paper | Landing | White | -2.129 | 1.266 | -1.681 | 0.212 |
|  | Pan Trap - Paper | Landing | White | 0.256 | 0.658 | 0.389 | 0.919 |
|  | Flower - Pan Trap | Landing | Yellow | -1.008 | 0.647 | -1.556 | 0.264 |
|  | Flower - Paper | Landing | Yellow | -0.879 | 0.652 | -1.347 | 0.369 |
|  | Pan Trap - Paper | Landing | Yellow | 0.128 | 0.474 | 0.270 | 0.960 |
| Small Insects | Flower - Pan Trap | Approach | Blue | 1.145 | 0.468 | 2.444 | **0.038** |
|  | Flower - Paper | Approach | Blue | 1.500 | 0.541 | 2.771 | **0.015** |
|  | Pan Trap - Paper | Approach | Blue | 0.354 | 0.592 | 0.598 | 0.820 |
|  | Flower - Pan Trap | Approach | White | -0.009 | 0.372 | -0.025 | 0.999 |
|  | Flower - Paper | Approach | White | 1.314 | 0.487 | 2.698 | **0.019** |
|  | Pan Trap - Paper | Approach | White | 1.324 | 0.490 | 2.698 | **0.019** |
|  | Flower - Pan Trap | Approach | Yellow | -0.010 | 0.348 | -0.03 | 0.999 |
|  | Flower - Paper | Approach | Yellow | 0.742 | 0.447 | 1.656 | 0.221 |
|  | Pan Trap - Paper | Approach | Yellow | 0.752 | 0.451 | 1.666 | 0.218 |
|  | Flower - Pan Trap | Landing | Blue | 1.107 | 0.341 | 3.240 | **0.003** |
|  | Flower - Paper | Landing | Blue | 1.037 | 0.338 | 3.062 | **0.006** |
|  | Pan Trap - Paper | Landing | Blue | -0.069 | 0.411 | -0.169 | 0.984 |
|  | Flower - Pan Trap | Landing | White | -0.047 | 0.181 | -0.261 | 0.962 |
|  | Flower - Paper | Landing | White | 0.852 | 0.235 | 3.612 | ***P < 0.001*** |
|  | Pan Trap - Paper | Landing | White | 0.899 | 0.233 | 3.859 | ***P < 0.001*** |
|  | Flower - Pan Trap | Landing | Yellow | -0.048 | 0.122 | -0.396 | 0.916 |
|  | Flower - Paper | Landing | Yellow | 0.279 | 0.133 | 2.100 | 0.089 |
|  | Pan Trap - Paper | Landing | Yellow | 0.328 | 0.132 | 2.483 | **0.034** |
| Other Flies | Flower - Pan Trap | Approach | Blue | 0.795 | 0.562 | 1.413 | 0.333 |
|  | Flower - Paper | Approach | Blue | 0.703 | 0.554 | 1.269 | 0.412 |
|  | Pan Trap - Paper | Approach | Blue | -0.091 | 0.574 | -0.158 | 0.986 |
|  | Flower - Pan Trap | Approach | White | 0.010 | 0.564 | 0.019 | 0.999 |
|  | Flower - Paper | Approach | White | -0.367 | 0.555 | -0.661 | 0.785 |
|  | Pan Trap - Paper | Approach | White | -0.378 | 0.514 | -0.735 | 0.742 |
|  | Flower - Pan Trap | Approach | Yellow | 0.249 | 0.423 | 0.589 | 0.825 |
|  | Flower - Paper | Approach | Yellow | 0.472 | 0.434 | 1.088 | 0.521 |
|  | Pan Trap - Paper | Approach | Yellow | 0.223 | 0.458 | 0.487 | 0.877 |
|  | Flower - Pan Trap | Landing | Blue | -0.316 | 0.444 | -0.712 | 0.755 |
|  | Flower - Paper | Landing | Blue | -0.480 | 0.430 | -1.115 | 0.504 |
|  | Pan Trap - Paper | Landing | Blue | -0.164 | 0.408 | -0.401 | 0.914 |
|  | Flower - Pan Trap | Landing | White | -1.100 | 0.396 | -2.772 | **0.015** |
|  | Flower - Paper | Landing | White | -1.552 | 0.377 | -4.113 | ***P < 0.001*** |
|  | Pan Trap - Paper | Landing | White | -0.451 | 0.256 | -1.757 | 0.183 |
|  | Flower - Pan Trap | Landing | Yellow | -0.861 | 0.254 | -3.387 | **0.002** |
|  | Flower - Paper | Landing | Yellow | -0.711 | 0.259 | -2.742 | **0.016** |
|  | Pan Trap - Paper | Landing | Yellow | 0.150 | 0.207 | 0.722 | 0.75 |

Table 6. **Individual insect group analysis of attractant preference when considering interactions with only behaviour (~attractant|behaviour)**

Results from pairwise test on emmeans post-hoc tests on negative binomial GLMMs of insect count data, contrasting the different attractants when considering their interactions with behaviour. Each insect group was analysed separately.

Values in bold signify that there is a significant behavioural response occurring.

P < 0.001 = high significance, P < 0.01 = medium significance, P < 0.05 = low significance.

| **Insect** | **Attractant Contrast** | **Behaviour** | **Estimate** | **SE** | **Z ratio** | **P value** |
| --- | --- | --- | --- | --- | --- | --- |
| Hoverflies | Flower – Pan Trap | Approach | 0.158 | 0.230 | 0.687 | 0.770 |
|  | Flower – Paper | Approach | 0.352 | 0.247 | 1.427 | 0.326 |
|  | Pan Trap – Paper | Approach | 0.194 | 0.262 | 0.737 | 0.740 |
|  | Flower – Pan Trap | Landing | 1.497 | 0.352 | 4.250 | ***P < 0.001*** |
|  | Flower – Paper | Landing | 2.437 | 0.506 | 4.812 | ***P < 0.001*** |
|  | Pan Trap – Paper | Landing | 0.939 | 0.561 | 1.674 | 0.214 |
| Bumblebees | Flower – Pan Trap | Approach | 0.786 | 0.35 | 2.247 | 0.063 |
|  | Flower – Paper | Approach | 0.808 | 0.336 | 2.400 | **0.043** |
|  | Pan Trap – Paper | Approach | 0.021 | 0.396 | 0.054 | 0.998 |
|  | Flower – Pan Trap | Landing | -0.040 | 1.025 | -0.039 | 0.999 |
|  | Flower – Paper | Landing | 22.134 | 38811.530 | 0.001 | 1.000 |
|  | Pan Trap – Paper | Landing | 22.175 | 38811.530 | 0.001 | 1.000 |
| Solitary Bees | Flower – Pan Trap | Approach | -0.329 | 0.349 | -0.943 | 0.612 |
|  | Flower – Paper | Approach | 0.152 | 0.407 | 0.374 | 0.925 |
|  | Pan Trap – Paper | Approach | 0.482 | 0.383 | 1.256 | 0.420 |
|  | Flower – Pan Trap | Landing | 0.032 | 0.432 | 0.074 | 0.996 |
|  | Flower – Paper | Landing | 0.856 | 0.529 | 1.618 | 0.237 |
|  | Pan Trap – Paper | Landing | 0.824 | 0.517 | 1.592 | 0.248 |
| Wasps | Flower – Pan Trap | Approach | -1.82 | 0.557 | -3.266 | **0.003** |
|  | Flower – Paper | Approach | -1.48 | 0.571 | -2.591 | **0.025** |
|  | Pan Trap – Paper | Approach | 0.339 | 0.313 | 1.084 | 0.523 |
|  | Flower – Pan Trap | Landing | -1.675 | 0.802 | -2.086 | 0.092 |
|  | Flower – Paper | Landing | -1.459 | 0.81 | -1.800 | 0.169 |
|  | Pan Trap – Paper | Landing | 0.215 | 0.515 | 0.418 | 0.907 |
| Small Insects | Flower – Pan Trap | Approach | 0.375 | 0.352 | 1.065 | 0.535 |
|  | Flower – Paper | Approach | 1.185 | 0.45 | 2.634 | **0.022** |
|  | Pan Trap – Paper | Approach | 0.81 | 0.46 | 1.759 | 0.183 |
|  | Flower – Pan Trap | Landing | 0.337 | 0.137 | 2.450 | **0.037** |
|  | Flower – Paper | Landing | 0.723 | 0.146 | 4.922 | ***P < 0.001*** |
|  | Pan Trap – Paper | Landing | 0.386 | 0.165 | 2.331 | 0.051 |
| Other Flies | Flower – Pan Trap | Approach | 0.351 | 0.427 | 0.822 | 0.688 |
|  | Flower – Paper | Approach | 0.269 | 0.43 | 0.627 | 0.805 |
|  | Pan Trap – Paper | Approach | -0.082 | 0.456 | -0.180 | 0.982 |
|  | Flower – Pan Trap | Landing | -0.759 | 0.226 | -3.359 | **0.002** |
|  | Flower – Paper | Landing | -0.914 | 0.22 | -4.152 | ***P < 0.001*** |
|  | Pan Trap – Paper | Landing | -0.155 | 0.179 | -0.863 | 0.663 |

Table 7. **Individual insect group analysis of colour preference when considering interactions with behaviour (~colour|behaviour)**

Results from a pairwise test on emmeans post-hoc tests on negative binomial GLMMs of insect count data, contrasting the different colour preferences when considering their interactions with behaviour. Each insect group was analysed separately.

Values in bold signify that there is a significant behavioural response occurring.

P < 0.001 = high significance, P < 0.01 = medium significance, P < 0.05 = low significance.

| **Insect** | **Colour Contrast** | **Behaviour** | **Estimate** | **SE** | **Z ratio** | **P value** |
| --- | --- | --- | --- | --- | --- | --- |
| Hoverflies | Blue - White | Approach | 0.211 | 0.304 | 0.695 | 0.765 |
|  | Blue - Yellow | Approach | -1.866 | 0.219 | -8.508 | ***P < 0.001*** |
|  | White - Yellow | Approach | -2.078 | 0.239 | -8.667 | ***P < 0.001*** |
|  | Blue - White | Landing | -0.650 | 0.613 | -1.061 | 0.538 |
|  | Blue - Yellow | Landing | -2.573 | 0.505 | -5.092 | ***P < 0.001*** |
|  | White - Yellow | Landing | -1.922 | 0.406 | -4.728 | ***P < 0.001*** |
| Bumblebees | Blue - White | Approach | 1.137 | 0.394 | 2.887 | **0.010** |
|  | Blue - Yellow | Approach | 0.202 | 0.299 | 0.675 | 0.777 |
|  | White - Yellow | Approach | -0.935 | 0.398 | -2.346 | **0.049** |
|  | Blue - White | Landing | -0.411 | 1.037 | -0.396 | 0.917 |
|  | Blue - Yellow | Landing | 18.413 | 7241.386 | 0.002 | 0.999 |
|  | White - Yellow | Landing | 18.825 | 7241.386 | 0.002 | 0.999 |
| Solitary Bees | Blue - White | Approach | 0.034 | 0.436 | 0.078 | 0.996 |
|  | Blue - Yellow | Approach | -0.923 | 0.358 | -2.577 | **0.026** |
|  | White - Yellow | Approach | -0.958 | 0.367 | -2.606 | **0.024** |
|  | Blue - White | Landing | -0.777 | 0.663 | -1.171 | 0.469 |
|  | Blue - Yellow | Landing | -1.921 | 0.592 | -3.245 | **0.003** |
|  | White - Yellow | Landing | -1.143 | 0.432 | -2.642 | **0.022** |
| Wasps | Blue - White | Approach | -0.566 | 0.569 | -0.996 | 0.579 |
|  | Blue - Yellow | Approach | -1.465 | 0.473 | -3.092 | **0.005** |
|  | White - Yellow | Approach | -0.898 | 0.428 | -2.094 | 0.090 |
|  | Blue - White | Landing | -0.207 | 0.990 | -0.209 | 0.976 |
|  | Blue - Yellow | Landing | -2.422 | 0.801 | -3.023 | **0.007** |
|  | White - Yellow | Landing | -2.215 | 0.713 | -3.105 | **0.005** |
| Small Insects | Blue - White | Approach | -0.904 | 0.550 | -1.643 | 0.227 |
|  | Blue - Yellow | Approach | -1.867 | 0.501 | -3.727 | ***P < 0.001*** |
|  | White - Yellow | Approach | -0.963 | 0.360 | -2.674 | **0.020** |
|  | Blue - White | Landing | -1.135 | 0.180 | -6.304 | ***P < 0.001*** |
|  | Blue - Yellow | Landing | -2.133 | 0.164 | -12.965 | ***P < 0.001*** |
|  | White - Yellow | Landing | -0.997 | 0.105 | -9.474 | ***P < 0.001*** |
| Other Flies | Blue - White | Approach | 0.285 | 0.594 | 0.480 | 0.880 |
|  | Blue - Yellow | Approach | -1.262 | 0.438 | -2.879 | **0.011** |
|  | White - Yellow | Approach | -1.548 | 0.497 | -3.111 | **0.005** |
|  | Blue - White | Landing | -0.711 | 0.239 | -2.972 | **0.008** |
|  | Blue - Yellow | Landing | -1.295 | 0.215 | -6.022 | ***P < 0.001*** |
|  | White - Yellow | Landing | -0.583 | 0.177 | -3.279 | **0.002** |

Table 8. **Hurdle model analysis of time spent landed on the attractants. Analysis of attractant spent longer on when considering interactions with insect groups and colour (~attractant|insect|colour)**

Results from a pairwise test on emmeans post-hoc test on hurdle model comparing landing times of all the relevant insect groups. This table contrasts attractants in regards to their interactions with colour and insect group. Values in bold signify that there is a significant behavioural response occurring.
P < 0.001 = high significance, P < 0.01 = medium significance, P < 0.05 = low significance.

| **Insect** | **Contrast** | **Colour** | **Estimate** | **SE** | **T ratio** | **P value** |
| --- | --- | --- | --- | --- | --- | --- |
| Beetles | Flower - Pan Trap | Blue | -26.516 | 33.876 | -0.782 | 0.713 |
|  | Flower - Paper | Blue | -5.983 | 4.419 | -1.354 | 0.365 |
|  | Pan Trap - Paper | Blue | 20.533 | 32.793 | 0.626 | 0.805 |
|  | Flower - Pan Trap | White | -16.226 | 13.966 | -1.161 | 0.476 |
|  | Flower - Paper | White | -2.431 | 2.503 | -0.971 | 0.595 |
|  | Pan Trap - Paper | White | 13.794 | 13.496 | 1.022 | 0.562 |
|  | Flower - Pan Trap | Yellow | -0.969 | 1.274 | -0.760 | 0.727 |
|  | Flower - Paper | Yellow | -0.176 | 0.207 | -0.853 | 0.669 |
|  | Pan Trap - Paper | Yellow | 0.793 | 1.165 | 0.680 | 0.775 |
| Hoverflies | Flower - Pan Trap | Blue | 0.492 | 0.300 | 1.640 | 0.228 |
|  | Flower - Paper | Blue | 0.527 | 0.322 | 1.636 | 0.230 |
|  | Pan Trap - Paper | Blue | 0.034 | 0.040 | 0.860 | 0.665 |
|  | Flower - Pan Trap | White | 1.465 | 0.900 | 1.627 | 0.234 |
|  | Flower - Paper | White | 1.974 | 1.058 | 1.866 | 0.148 |
|  | Pan Trap - Paper | White | 0.508 | 0.451 | 1.128 | 0.496 |
|  | Flower - Pan Trap | Yellow | 4.576 | 4.677 | 0.978 | 0.590 |
|  | Flower - Paper | Yellow | 11.924 | 3.205 | 3.720 | **P < 0.001** |
|  | Pan Trap - Paper | Yellow | 7.348 | 3.916 | 1.876 | 0.145 |
| Other Flies | Flower - Pan Trap | Blue | -3.862 | 2.137 | -1.807 | 0.167 |
|  | Flower - Paper | Blue | -9.636 | 3.925 | -2.454 | **0.037** |
|  | Pan Trap - Paper | Blue | -5.773 | 4.164 | -1.386 | 0.348 |
|  | Flower - Pan Trap | White | -17.404 | 5.172 | -3.364 | **0.002** |
|  | Flower - Paper | White | -24.326 | 6.195 | -3.926 | **P < 0.001** |
|  | Pan Trap - Paper | White | -6.922 | 7.240 | -0.956 | 0.604 |
|  | Flower - Pan Trap | Yellow | -16.928 | 4.931 | -3.432 | **0.001** |
|  | Flower - Paper | Yellow | -26.304 | 6.841 | -3.844 | **P < 0.001** |
|  | Pan Trap - Paper | Yellow | -9.376 | 7.434 | -1.261 | 0.417 |
| Other Insects | Flower - Pan Trap | Blue | 10.564 | 6.772 | 1.559 | 0.263 |
|  | Flower - Paper | Blue | 14.463 | 6.909 | 2.093 | 0.091 |
|  | Pan Trap - Paper | Blue | 3.898 | 2.952 | 1.320 | 0.383 |
|  | Flower - Pan Trap | White | 6.444 | 16.576 | 0.388 | 0.920 |
|  | Flower - Paper | White | 24.192 | 19.205 | 1.259 | 0.418 |
|  | Pan Trap - Paper | White | 17.748 | 12.450 | 1.425 | 0.327 |
|  | Flower - Pan Trap | Yellow | 27.648 | 41.262 | 0.670 | 0.780 |
|  | Flower - Paper | Yellow | 73.326 | 37.874 | 1.936 | 0.128 |
|  | Pan Trap - Paper | Yellow | 45.677 | 25.286 | 1.806 | 0.167 |
| Small Insects | Flower - Pan Trap | Blue | 8.564 | 9.552 | 0.896 | 0.642 |
|  | Flower - Paper | Blue | 7.753 | 9.886 | 0.784 | 0.712 |
|  | Pan Trap - Paper | Blue | -0.811 | 9.661 | -0.083 | 0.996 |
|  | Flower - Pan Trap | White | -2.877 | 8.711 | -0.330 | 0.941 |
|  | Flower - Paper | White | -4.161 | 11.534 | -0.360 | 0.930 |
|  | Pan Trap - Paper | White | -1.283 | 11.320 | -0.113 | 0.992 |
|  | Flower - Pan Trap | Yellow | -3.921 | 10.643 | -0.368 | 0.927 |
|  | Flower - Paper | Yellow | -27.111 | 14.389 | -1.884 | 0.143 |
|  | Pan Trap - Paper | Yellow | -23.189 | 14.327 | -1.618 | 0.237 |
| Solitary Bees | Flower - Pan Trap | Blue | 0.372 | 0.339 | 1.096 | 0.516 |
|  | Flower - Paper | Blue | 0.361 | 0.3400 | 1.062 | 0.537 |
|  | Pan Trap - Paper | Blue | -0.011 | 0.149 | -0.077 | 0.996 |
|  | Flower - Pan Trap | White | 1.134 | 3.817 | 0.297 | 0.952 |
|  | Flower - Paper | White | 2.372 | 3.708 | 0.639 | 0.798 |
|  | Pan Trap - Paper | White | 1.237 | 2.771 | 0.446 | 0.895 |
|  | Flower - Pan Trap | Yellow | 2.463 | 4.136 | 0.595 | 0.822 |
|  | Flower - Paper | Yellow | 3.582 | 4.274 | 0.838 | 0.679 |
|  | Pan Trap - Paper | Yellow | 1.118 | 3.139 | 0.356 | 0.932 |
| Wasps | Flower - Pan Trap | Blue | -0.404 | 0.525 | -0.770 | 0.721 |
|  | Flower - Paper | Blue | -0.515 | 0.727 | -0.708 | 0.758 |
|  | Pan Trap - Paper | Blue | -0.111 | 0.495 | -0.224 | 0.972 |
|  | Flower - Pan Trap | White | -1.080 | 1.230 | -0.878 | 0.653 |
|  | Flower - Paper | White | -0.803 | 0.798 | -1.005 | 0.573 |
|  | Pan Trap - Paper | White | 0.277 | 0.897 | 0.308 | 0.948 |
|  | Flower - Pan Trap | Yellow | -6.256 | 2.846 | -2.198 | 0.071 |
|  | Flower - Paper | Yellow | -5.028 | 2.898 | -1.734 | 0.192 |
|  | Pan Trap - Paper | Yellow | 1.228 | 3.975 | 0.309 | 0.948 |

Table 9. **Hurdle model analysis of time spent landed on the attractants. Analysis of attractant spent longer on when considering interactions with insect group(~attractant|insect)**

Results from a pairwise test on emmeans post-hoc test on hurdle model comparing landing times of all the relevant insect groups. This table contrasts attractants in regards to their interactions with each insect group. Values in bold signify that there is a significant behavioural response occurring.
P < 0.001 = high significance, P < 0.01 = medium significance, P < 0.05 = low significance.

| **Insect** | **Attractant Contrast** | **Estimate** | **SE** | **T ratio** | **P value** |
| --- | --- | --- | --- | --- | --- |
| Beetles | Flower - Pan Trap | -14.57 | 13.814 | -1.054 | 0.542 |
|  | Flower - Paper | -2.863 | 1.864 | -1.536 | 0.274 |
|  | Pan Trap - Paper | 11.707 | 13.923 | 0.840 | 0.677 |
| Hoverflies | Flower - Pan Trap | 2.178 | 1.713 | 1.271 | 0.411 |
|  | Flower - Paper | 4.809 | 1.167 | 4.120 | **P < 0.001** |
|  | Pan Trap - Paper | 2.630 | 1.399 | 1.880 | 0.144 |
| Other Flies | Flower - Pan Trap | -12.731 | 3.169 | -4.017 | **P < 0.001** |
|  | Flower - Paper | -20.089 | 4.076 | -4.928 | **P < 0.001** |
|  | Pan Trap - Paper | -7.357 | 4.788 | -1.536 | 0.273 |
| Other Insects | Flower - Pan Trap | 14.885 | 19.734 | 0.754 | 0.731 |
|  | Flower - Paper | 37.327 | 17.395 | 2.145 | 0.081 |
|  | Pan Trap - Paper | 22.441 | 11.261 | 1.992 | 0.114 |
| Small Insects | Flower - Pan Trap | 0.588 | 6.083 | 0.096 | 0.994 |
|  | Flower - Paper | -7.839 | 7.683 | -1.020 | 0.564 |
|  | Pan Trap - Paper | -8.428 | 7.920 | -1.064 | 0.536 |
| Solitary Bees | Flower - Pan Trap | 1.323 | 2.575 | 0.513 | 0.864 |
|  | Flower - Paper | 2.105 | 2.548 | 0.826 | 0.686 |
|  | Pan Trap - Paper | 0.781 | 1.929 | 0.405 | 0.913 |
| Wasps | Flower - Pan Trap | -2.580 | 1.202 | -2.145 | 0.081 |
|  | Flower - Paper | -2.115 | 1.159 | -1.824 | 0.161 |
|  | Pan Trap - Paper | 0.464 | 1.713 | 0.271 | 0.960 |

Table 10. **Hurdle model analysis of time spent landed on the attractants. Analysis of colour spent longer on when considering interactions with insect group (~colour|insect)**

Results from a pairwise test on emmeans post-hoc test on hurdle model comparing landing times of all the relevant insect groups. This table contrasts colours in regards to their interactions with each insect group. Values in bold signify that there is a significant behavioural response occurring.
P < 0.001 = high significance, P < 0.01 = medium significance, P < 0.05 = low significance.

| **Insect** | **Colour Contrast** | **Estimate** | **SE** | **T ratio** | **P value** |
| --- | --- | --- | --- | --- | --- |
| Beetles | Blue - White | 4.961 | 12.155 | 0.408 | 0.912 |
|  | Blue - Yellow | 10.894 | 12.063 | 0.903 | 0.638 |
|  | White - Yellow | 5.932 | 4.932 | 1.202 | 0.451 |
| Hoverflies | Blue - White | -0.722 | 0.514 | -1.402 | 0.339 |
|  | Blue - Yellow | -8.037 | 1.677 | -4.792 | **P < 0.001** |
|  | White - Yellow | -7.315 | 1.730 | -4.227 | **P < 0.001** |
| Other Flies | Blue - White | -12.446 | 3.843 | -3.238 | **0.003** |
|  | Blue - Yellow | -16.465 | 3.676 | -4.478 | **P < 0.001** |
|  | White - Yellow | -4.018 | 4.547 | -0.883 | 0.65 |
| Other Insects | Blue - White | -9.484 | 10.124 | -0.936 | 0.617 |
|  | Blue - Yellow | -39.725 | 17.895 | -2.219 | 0.068 |
|  | White - Yellow | -30.240 | 19.396 | -1.559 | 0.263 |
| Small Insects | Blue - White | -20.088 | 6.904 | -2.909 | **0.010** |
|  | Blue - Yellow | -60.575 | 7.528 | -8.045 | **P < 0.001** |
|  | White - Yellow | -40.487 | 7.413 | -5.461 | **P < 0.001** |
| Solitary Bees | Blue - White | -3.841 | 2.625 | -1.463 | 0.308 |
|  | Blue - Yellow | -5.031 | 1.811 | -2.776 | **0.015** |
|  | White - Yellow | -1.189 | 3.150 | -0.377 | 0.924 |
| Wasps | Blue - White | -0.309 | 0.829 | -0.372 | 0.926 |
|  | Blue - Yellow | -3.899 | 1.426 | -2.734 | **0.017** |
|  | White - Yellow | -3.590 | 1.558 | -2.303 | 0.055 |
